# Co-option of lysosomal machinery shapes the symbiosis supporting coral reefs

**DOI:** 10.1101/2025.10.09.679812

**Authors:** Shumpei Maruyama, Catherine F. Henderson, Natalie Swinhoe, Griffin P. Kowalewski, Emily K. Meier, Ty R. Engelke, Phillip A. Cleves

## Abstract

Intracellular photosymbiosis has evolved across life and forms the foundation of coral reef ecosystems. Using the sea anemone Aiptasia as a model, we generated a high-quality proteome of the symbiosome, the organelle that houses algal symbionts. This proteome revealed protein trafficking mechanisms and the types of biomolecules exchanged during symbiosis. Symbiosomal enrichment of lysosomal proteins, visualization of lysosomal fusion, along with reduced symbiosis following knockdown of lysosomal genes, supports its phagolysosomal identity and that extensive co-option of lysosomal proteins shapes the symbiosome. CRISPR/Cas9-induced mutations in the symbiosomal and lysosomal bicarbonate/sulfate transporter, SLC26A11, disrupted symbiosis in both Aiptasia and a reef-building coral. These findings support that anemones and corals independently evolved a carbon-concentrating and sulfate transport mechanism to fuel photosymbiosis by co-opting an orthologous lysosomal transporter.

## Main Text

Intracellular photosymbiosis has played a key role in the evolution of eukaryotic cellular complexity and has repeatedly evolved across life, allowing taxa to expand their ecological niches (*1*, *2*). Nutritional exchange between symbiotic partners is a key feature of photosymbiotic relationships that have shaped the evolution of organisms in each eukaryotic kingdom [*e.g.,* chloroplasts in plants and protists (*2*), lichen-algal symbioses in fungi (*3*), and coral-algal symbiosis in animals (*1*)]. In reef-building corals, photosymbiosis has evolved multiple times (*4*) [as well as in other cnidarians (*5*)], which fuels the energetic requirements of coral reefs. This nutritional symbiosis between corals and dinoflagellate algae (family Symbiodiniaceae) allows corals to thrive by the provision of photosynthetic products from the algae (*6*). In return, the host provides the algae with inorganic carbon and nutrients required for photosynthesis and algal physiology (*7*).

Most coral species acquire algal symbionts each generation from the environment, where the algae are phagocytosed into specialized gastrodermal cells (called ‘symbiocytes’), processed through the endomembrane trafficking system, and then maintained in a specialized organelle called the symbiosome (*7*). The host-derived symbiosome membrane serves as the cellular interface between the two partners. This membrane is the site of interpartner communication and nutrient exchange and defines the local environment for the alga (*8*). The symbiosome is an acidic organelle (pH ∼ 4) (*9*), shown to translocate a variety of metabolites [*e.g.*, glucose (*10*), sterols (*11*), ammonium (*12*, *13*), and other lipids (*14*)]. Over the past several decades, research on corals and an anemone model for coral symbiosis, Aiptasia (*sensu stricto Exaiptasia diaphana*), has attempted to characterize the symbiosome proteome to better understand the symbiosis and the proteins that mediate organelle biogenesis and metabolite transport (*15*). A previous study sought to generate a symbiosome proteome in Aiptasia but detected only 17 proteins, of which many appear non-specific (*e.g.*, actin, heat-shock proteins) (*16*). Due to the difficulties of purifying the symbiosome membrane and proteomics, research has focused on immunofluorescence assays, which have identified several transporters for glucose (*12*, *17*), sterols (*11*, *18*), and ammonium (*12*, *13*, *17*), that localize to the symbiosome, corroborating earlier functional studies indicating that these metabolites are exchanged across the symbiosome. In addition, several markers of various endomembrane compartments have been localized to the symbiosome [*e.g.*, RAB5B (*19*), and phagolysosomal markers, V-ATPase (*9*), LAMP1A (*20*), RAB7A (*21*, *22*), mTOR (*23*)] resulting in controversy about whether the symbiosome is an arrested phagosome or a phagolysosome (*19*, *21*, *24–26*).

In this work, we perform proteomics on the symbiosome in Aiptasia and functionally characterized symbiosomal genes in both Aiptasia and a reef-building coral, *Galaxea fascicularis*. These results revealed a more conclusive understanding of the cellular origins of the symbiosome, how proteins are trafficked to the symbiosome, and the identification of other biomolecules exchanged across the membrane during symbiosis. We found that the symbiosome is formed through extensive co-option of lysosomal machinery, which are needed for symbiosis. These results provide an explanation for the repeated evolution of photosymbiosis in cnidarians and insights into the molecular basis of coral reef productivity.

### Biochemical purification of symbiosomes

To generate a high-quality symbiosome proteome, we developed a method to enrich and purify the symbiosome from adult Aiptasia. First, we developed an animal dissociation method, which results in a heterogenous mixture of cells, including symbiocytes, released symbiosome organelles containing algae, and free algae (Fig. 1A). We then developed a staining strategy to differentiate between symbiocytes, released symbiosomes, and free algae from the mixed population of dissociated cells using Calcein Violet to stain host cytoplasm and an antibody for a symbiosomal ammonium transporter, RHBG, to specifically label released symbiosomes (Fig. 1, B and C, fig. S1). Super-resolution confocal microscopy of released symbiosomes confirmed that RHBG stained only the outer symbiosome membrane (Fig. 1, D and E, fig. S2). We used the staining method to track the symbiosome membrane during a series of purification steps to isolate the symbiosome (Fig. 1, F to H; see materials and methods). We found that ∼80% of algae were in symbiosomes at the final step of the protocol before removal of the membrane, and ∼90% of the symbiosome membranes were removed after treatment with detergent (Fig. 1H). The supernatant, containing symbiosome membrane and contents, along with control samples from crude gastrodermal cell lysates (Fig. 1F, step 1) were then processed for proteomic analysis.

**Fig. 1.**
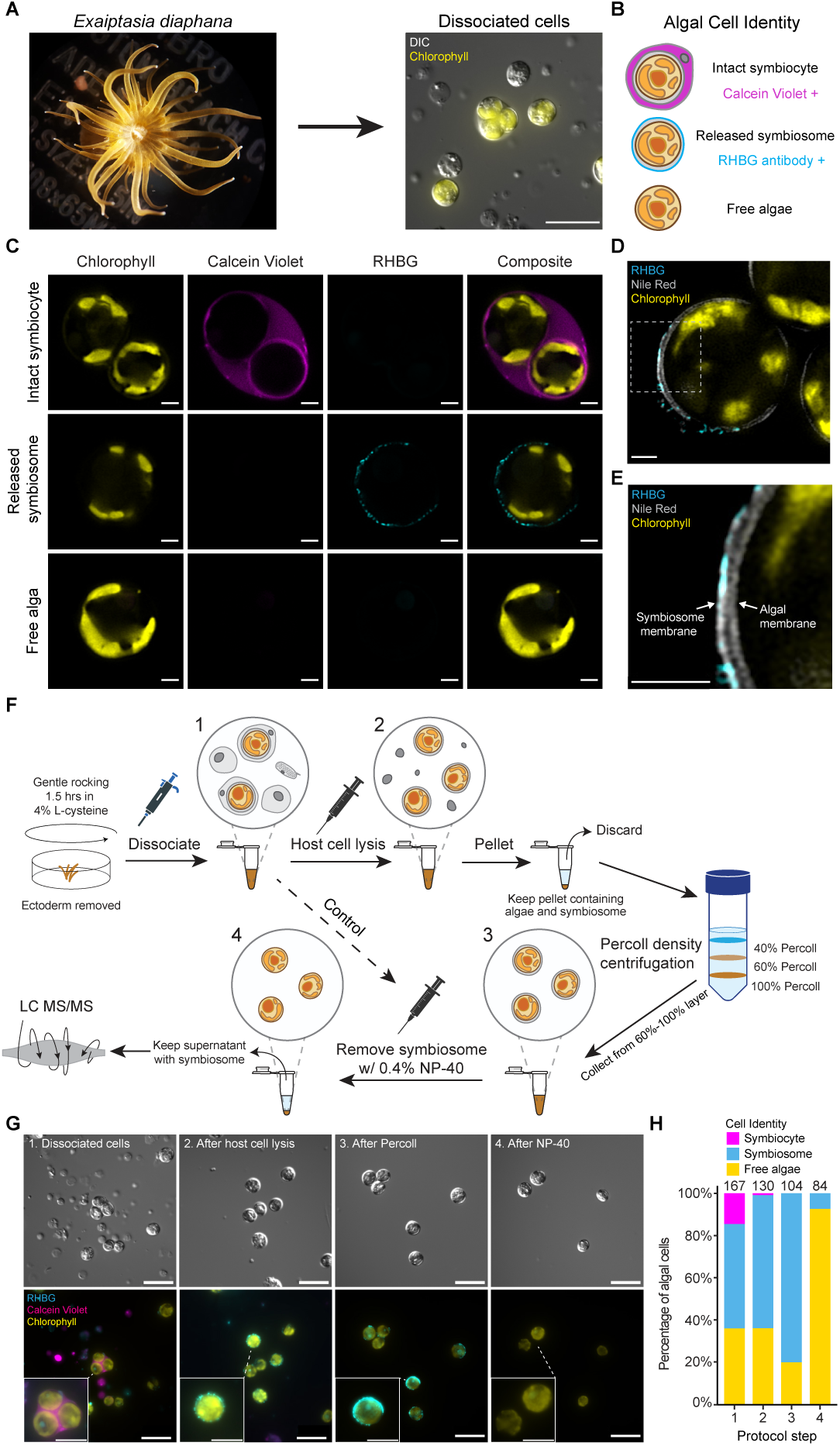
Purification of symbiosomes for proteomics. **(A)** Adult Aiptasia (left) and after dissociation into single cells (right). Dissociated cells include symbiont-containing cells (symbiocytes), released symbiosomes containing algae, and free algae. Algae can be identified by chlorophyll autofluorescence (yellow). 20 μm scale bar. **(B)** Staining strategy to distinguish symbiocytes, released symbiosomes, and free algae. The cytoplasmic stain (purple) stains only intact symbiocytes. An antibody for RHBG (cyan) stains only released symbiosomes, 2 μm scale bars. **(C)** Representative super-resolution confocal images of a stained symbiocyte, released symbiosome, and free alga. 2 μm scale bars. **(D)** Super-resolution confocal slice of a released symbiosome stained for RHBG protein and membrane (grey). 2 μm scale bar. **(E)** Higher magnification of white box in D allows for visualization of both the symbiosome and algal membranes. 2 μm scale bar. **(F)** Steps to purify symbiosomes (see materials and methods). **(G)** Representative differential interference contrast (top) and epifluorescence (bottom) images of cells stained at each step (1–4) of the protocol. 20 μm scale bars in larger images, 10 μm scale bars in insets. **(H)** Quantification of the proportion of symbiocytes, released symbiosomes, and free algae from a representative purification.

### Membrane trafficking, lysosomal, and transport proteins on the symbiosome membrane

To find proteins that localized to the symbiosome membrane, we identified proteins that were enriched in the symbiosome proteome compared to the control gastroderm proteome (Fig. 1F). Principal component analysis found that the majority of the variance (67.9%) was explained by the differences between symbiosome and control samples (Fig. 2A). We detected the enrichment of 200 proteins in the symbiosome, of which 49 were unique to the symbiosome samples (Fig. 2B, and table S1). We detected 19 proteins that are known to be on symbiosomes in our proteomics, of which 17 were enriched in the symbiosome samples (Fig. 2B, table S2) (*27*). As expected, we found a significant enrichment of transmembrane proteins on the symbiosome compared to the genome-wide proteome and control-enriched proteins, consistent with successful purification of the symbiosome membrane (Fig. 2C). There was also higher enrichment of proteins on the symbiosome that were transcriptionally upregulated in symbiotic relative to non-symbiotic anemones (Fig. 2D) (*27*). Interestingly, 80.5% of the symbiosome-enriched proteins were not transcriptionally upregulated in symbiotic animals suggesting that most of the symbiosome machinery have other roles in homeostasis and have been co-opted to function on the symbiosome.

**Fig. 2.**
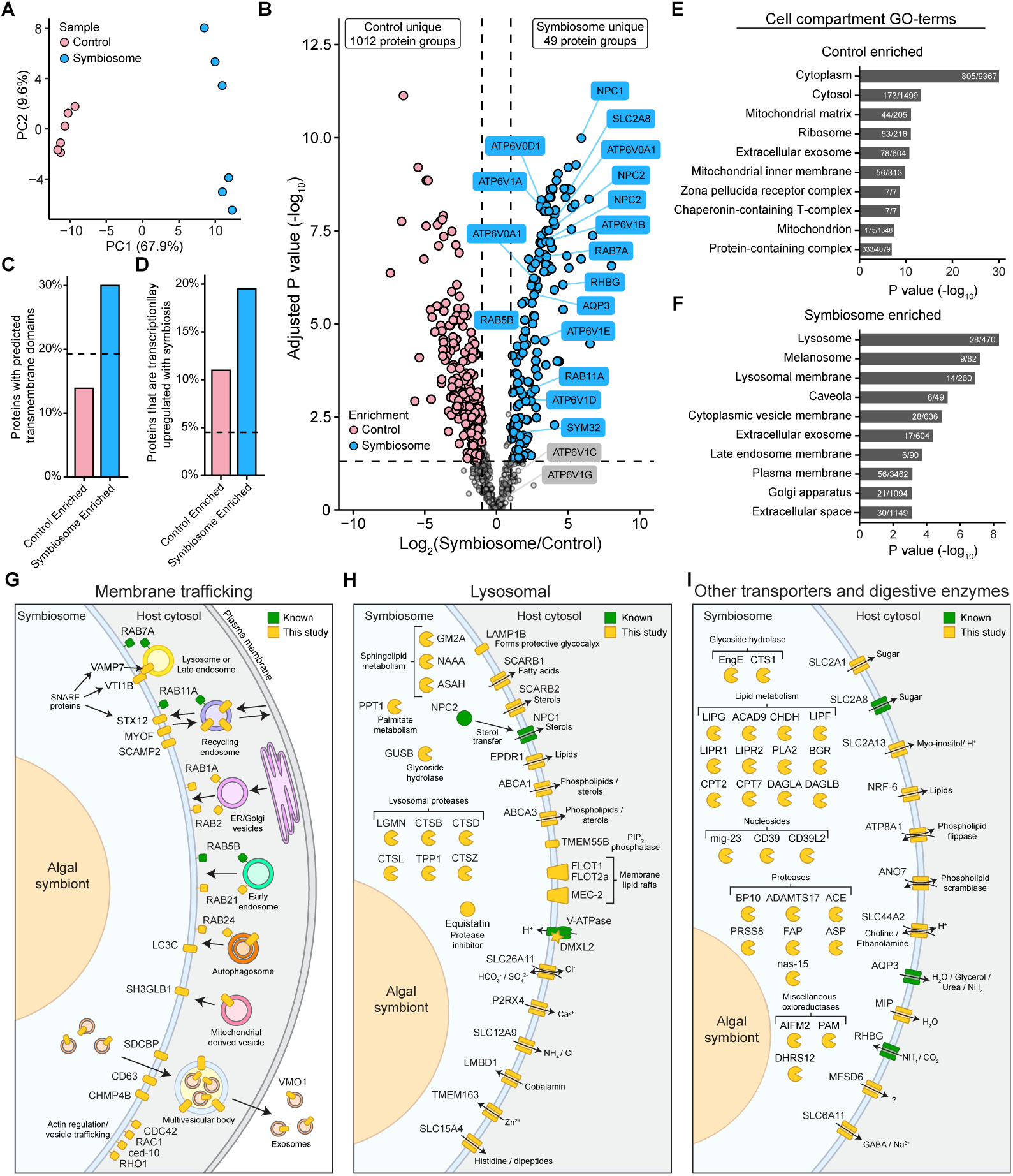
The symbiosome proteome. (**A**) Principal component analysis of control and symbiosome samples. **(B)** Differential enrichment of proteins between control and symbiosome samples. Annotated proteins are those previously localized to the symbiosome based on antibody staining in at least one symbiotic cnidarian (table S2). **(C)** Symbiosomal proteins are significantly enriched for transmembrane proteins compared to the genome-wide proteome (dashed line) or control-enriched proteins based on Chi-square tests (*P_adj_ =* 1.1 x 10^-16^ and *P_adj_* = 9.2 x 10^-29^, respectively). **(D)** Symbiosomal proteins are significantly enriched for genes that are transcriptionally upregulated by symbiosis (*27*) compared to the genome-wide proteome (dashed line) or control-enriched proteins based on Chi-square tests (*P_adj_ =* 1.7 x 10^-23^ and *P_adj_* = 6.7 x 10^-4^, respectively). **(E-F)** The top ten most significantly enriched cell-compartment gene-ontology (GO) terms for each group of proteins. White ratios indicate number of proteins found in the group over the number of proteins genome-wide that have the given GO-term. **(G-I)** Symbiosome proteins grouped by inferred function of orthologous genes from other organisms. Complete symbiosome protein list and citations for each predicted function are provided in table S1.

Cell compartment Gene Ontology (GO) analyses revealed the control-enriched proteins were associated with non-endosomal cellular compartments such as cytoplasm, mitochondria, and ribosomes (Fig. 2E and table S4), whereas the symbiosome-enriched proteins were associated with endomembrane organelles, including lysosomes, melanosomes, and late endosomes (Fig. 2F and table S4). Next, we predicted the function of each protein on the symbiosome based on their orthologous functions in other organisms and grouped them into four broad categories (Fig. 2, G to I and table S1). First, we identified a group of proteins involved in membrane trafficking (Fig. 2G and table S1). These proteins are predicted to orchestrate diverse trafficking mechanisms, including those involved in docking and fusing of vesicles, the discrimination of vesicle types (*e.g.*, recycling, early, and late endosomes and ER/Golgi vesicles), and actin cytoskeleton organization. Second, we detected a diversity of lysosomal proteins (Fig. 2H and table S1). Interestingly, these proteins include canonical proteins that mediate lysosome formation and function, including acidification machinery (*e.g.*, V-ATPase), lysosomal integral membrane proteins (*e.g.*, LAMP1B), and various lysosomal proteases (*e.g.*, cathepsins: CTSB, CTSD, CTSZ, and CTSL). For the cathepsins, we only detected peptides from the mature forms of each protein (fig. S3 and table S5), suggesting these are processed and active proteases (*28*). Third, the symbiosome contained a group of non-lysosomal digestive proteins predicted to break down carbohydrates, lipids, nucleosides, and proteins, indicating the symbiosome has robust digestive functions (Fig. 2I and table S1). Finally, we detected a variety of previously unknown membrane transport proteins that are predicted to shuttle metabolites known to cross the symbiosome [*e.g.*, sugars (*10*), sterols (*11*), lipids (*14*), ammonium (*12*, *13*)] and ions and metabolites not previously known to be transported (*e.g.*, calcium, bicarbonate, myo-inositol, histidine) (Fig. 2I and table S1). The myriad of digestive enzymes and transporters found in the symbiosome indicates a wide diversity of unappreciated metabolic pathways that integrate host and symbiont metabolisms.

### Dynamic membrane trafficking and phagosome-lysosome fusion shape the symbiosome

The presence of diverse membrane trafficking and lysosomal proteins suggests that the symbiosome forms from phagosome-lysosome fusion rather than phagosome arrest. To test this hypothesis, we visualized vesicle trafficking with membrane dyes and an antibody for a lysosomal marker LAMP1A, which forms a protective lysosomal glycocalyx and is known to be on the symbiosome (*20*) (Fig. 3A-C and fig. S4 and S5). We found that within symbiocytes, 95% of symbiosomes had at least one vesicle (Fig 3D), where 56.6% of these vesicles were LAMP1A-positive lysosomes with a mean diameter of 0.6 μm (Fig. 3E). The LAMP1A-negative vesicles with varying diameters suggest other vesicles are trafficked to and from the symbiosome, consistent with the diversity of trafficking proteins found in the proteome (Fig. 2G). Purification of the symbiosome significantly depleted smaller LAMP1A-positive lysosomes, indicating that other non-lysosomal vesicles remain tightly associated with the symbiosome (Fig. 3E). Consistently, we identified SNARE proteins in the symbiosome proteome that we predict are used to dock and tightly tether vesicles to the symbiosome (Fig. 2G and table S1). Together, these data indicate that lysosomes and at least one other type of vesicle dynamically fuse with the symbiosome, helping to shape the organelle.

**Fig. 3.**
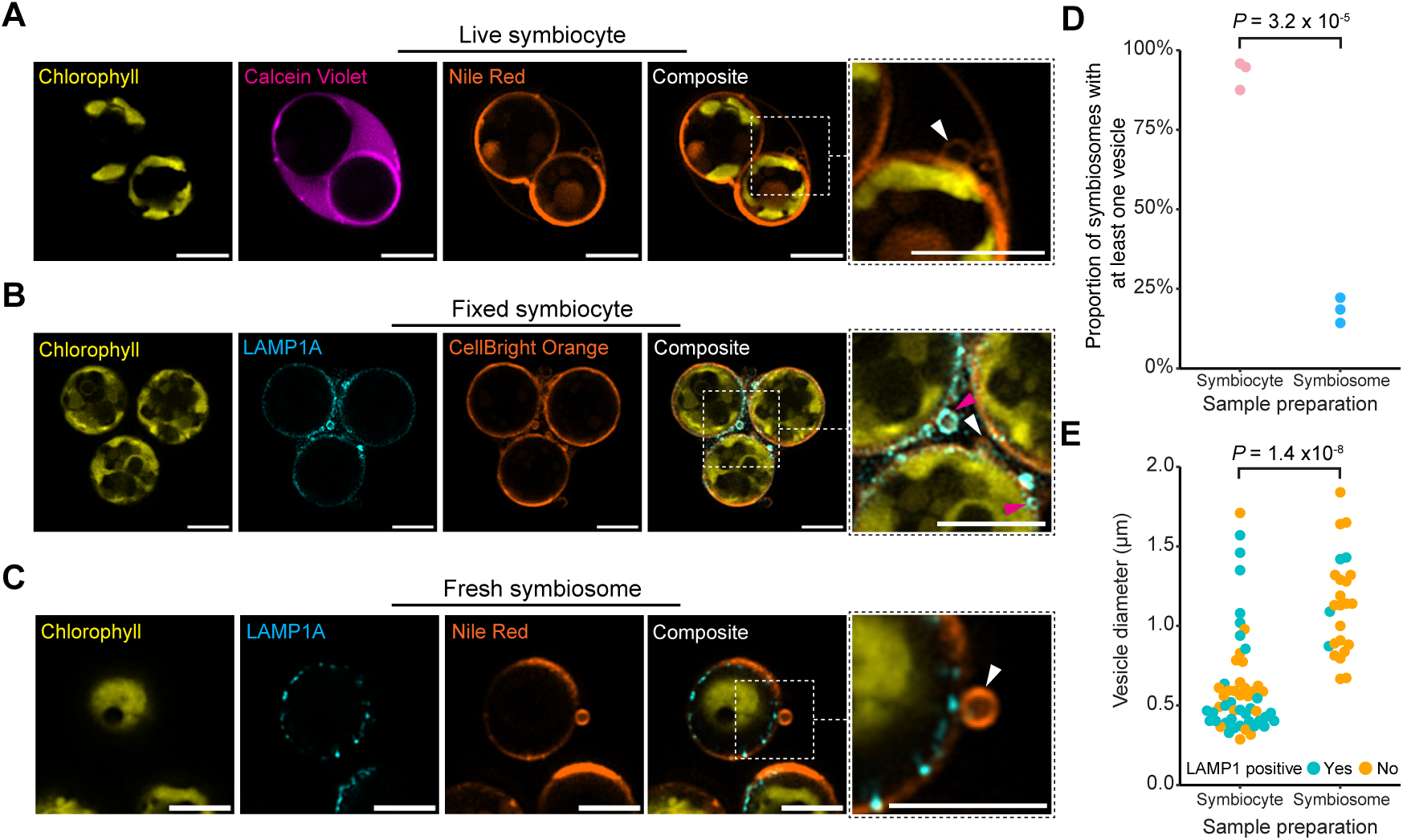
Lysosomes and other vesicles fuse to the symbiosome. **(A)** Live symbiocytes stained with a cytoplasmic dye (purple) and a membrane dye (orange). Nile red staining reveals vesicles tightly associate to the symbiosome membrane (white arrowhead). **(B)** Immunofluorescence of fixed symbiocytes reveals some vesicles are LAMP1A positive (magenta arrowhead) and LAMP1A negative (white arrowhead). **(C)** Immunofluorescence of live, purified symbiosomes show that vesicles are also found attached to freshly isolated symbiosomes (white arrowhead). 5 μm scales bars for (A-C). **(D)** Proportion of symbiosomes associated with vesicles in fixed symbiocytes and freshly isolated symbiosomes. *P* value determined by a Welch’s t-test. **(E)** Quantification of vesicles on fixed symbiocytes and freshly isolated symbiosomes. Each dot is a vesicle and *P* value determined by a Welch’s t-test.

### Co-option of phagolysosomal machinery is required for symbiosis

Because the fusion of lysosomes onto phagosomes is a terminal stage of phagolysosomal maturation (*29*), we hypothesized that the symbiosome has properties of a phagolysosome. Previous research has suggested that the symbiosomes of reef-building corals have a low pH from the action of V-ATPase (*9*). We found that the symbiosome is indeed acidic in Aiptasia by staining whole polyps with Lysotracker Green which stains acidic compartments (Fig. 4A and fig. S6). To evaluate the lysosomal functions of symbiosomes, we generated specific antibodies for two lysosomal proteins that were enriched in our symbiosomal proteome: LAMP1B, an ortholog to LAMP1A, and CTSB (fig. S4, S5, and S7). Using super-resolution confocal microscopy, we found that LAMP1A and LAMP1B were localized to the symbiosome membrane, and CTSB was localized to the symbiosome lumen (Fig. 4, B and C, and fig. S8). We also detected localization of LAMP1B and CTSB on small vesicles associated with the symbiosome membrane (fig. S9).

**Fig. 4.**
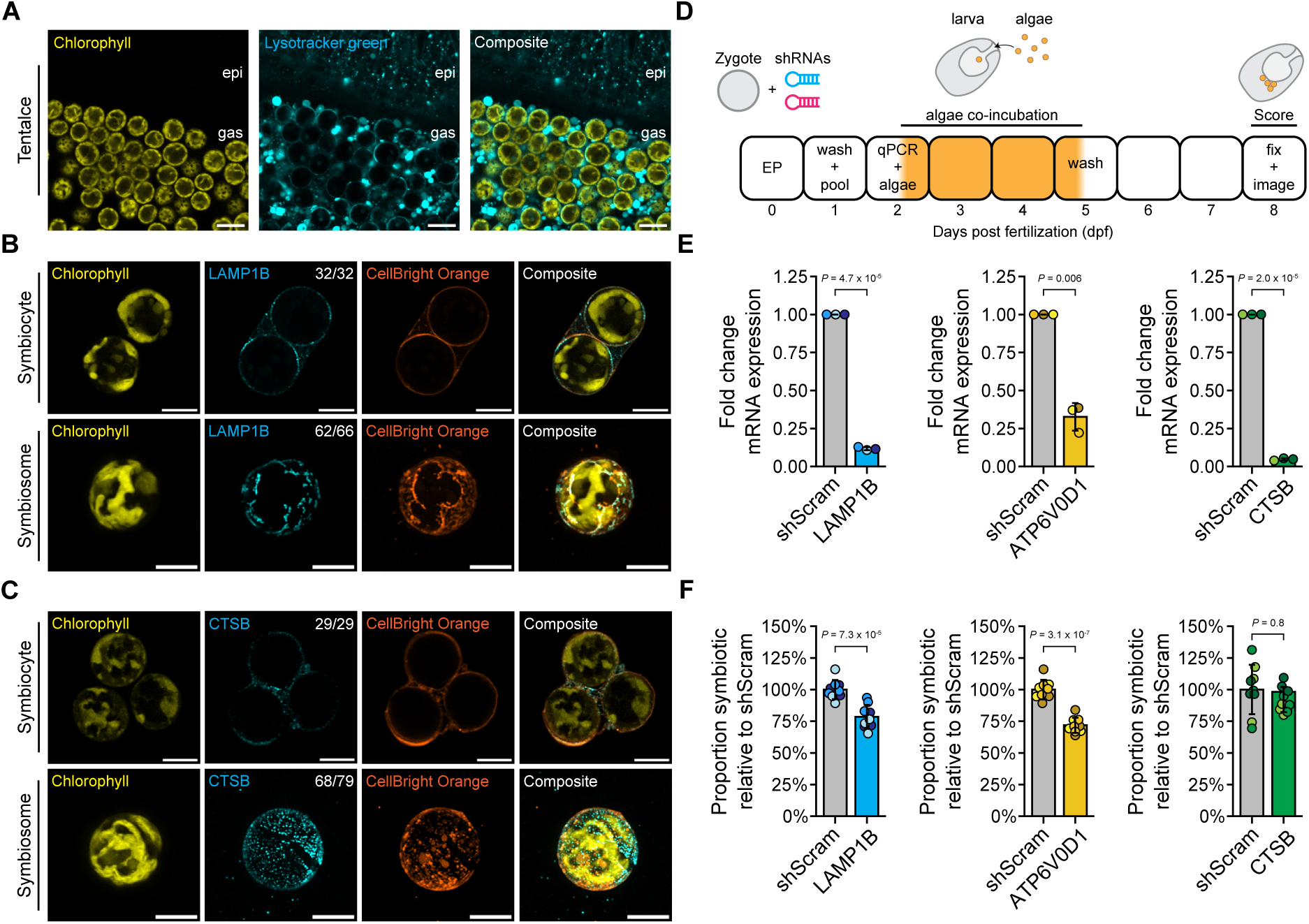
Knock down of lysosomal proteins on the symbiosome reduces symbiosis. **(A)** Live staining of whole animals with Lysotracker green shows that the symbiosome has an acidic pH consistent with the presence of V-ATPase subunits in the symbiosome proteome (Fig. 2H). 10 μm scale bars. **(B-C)** Immunofluorescent images of (B) LAMP1B and (C) CTSB showing localization to the symbiosome in symbiocytes and isolated symbiosomes. Numbers indicate fraction of cells with symbiosomal localization. 5 μm scale bars. **(D)** Strategy for RNAi experiments in Aiptasia larvae. **(E)** Efficient knock down of LAMP1B, ATP6V0D1, and CTSB mRNA levels at 2 dpf. qPCR results for three independent RNAi experiments are shown. **(F)** Knock down of LAMP1B and ATP6V0D1, but not CTSB reduce symbiosis after 6 days post inoculation with symbionts. The proportion of symbiotic larvae is shown for three biological replicates (∼100 larvae each) from each of three RNAi experiments. The different shades of color denote experimental replicates in **(E)** and **(F)**. *P* values were determined by Welch’s t-tests.

Our previous work found that knockdown of LAMP1A reduces symbiosis formation in Aiptasia larvae (*30*). We asked if the knockdown of other key lysosomal proteins [LAMP1B; CTSB; and a subunit of V-ATPase, ATP6V0D1] also impacted the formation of symbiosis. We knocked down each gene with shRNAs in Aiptasia zygotes, then quantified how many larvae became symbiotic after incubation with symbionts (Fig. 4D). We were able to successfully knock down mRNA expression of LAMP1B, ATP6V0D1, CTSB by 88%, 67%, and 95%, respectively (Fig. 4E). Knockdown of LAMP1B and ATP6V0D1 significantly reduced symbiosis formation (Fig. 4F). Conversely, knockdown of CTSB did not impact symbiosis formation (Fig. 4F), potentially due to redundancy in function by other cathepsins detected in the symbiosome proteome (Fig. 2H). Together, these observations support the phagolysosome-like function of symbiosomes and that lysosomal structural proteins and acidification machinery are important for forming symbiosis in Aiptasia.

### A symbiosomal bicarbonate/sulfate transporter is required for photosymbiosis in both Aiptasia and the coral, *Galaxea fascicularis*

A central question regarding photosymbiosis is how the host concentrates inorganic carbon and transports other essential metabolites (*e.g.*, sulfate) into the symbiosome to support algal growth and photosynthesis (*7*, *31*, *32*). For carbon concentration, it has been hypothesized that the action of an undescribed bicarbonate transporter on the symbiosome is required to concentrate the inorganic carbon needed to support the high rates of algal photosynthesis (*9*, *31*, *32*). In addition, the algae require a source of sulfate for amino acid synthesis and other key processes in the cell (*33*, *34*). However, the host transport mechanisms remain unclear. In our symbiosomal proteome, we identified a single candidate bicarbonate/sulfate transporter, SLC26A11. Orthologs of SLC26A11 have been shown to transport both bicarbonate and sulfate (*35*, *36*) driven by chloride exchange in various model systems (*35*), and the human ortholog has been localized to lysosomes (*37*).

To test if SLC26A11 is the symbiosomal transporter that could be supplying inorganic carbon and sulfate to the algae, we used a custom antibody for SLC26A11 to confirm its localization on the symbiosome membrane (Fig. 5A and fig. S4, S5, and S7). We found that SLC26A11, like LAMP1A, localized to small vesicles, consistent with its lysosomal localization (fig. S9). To determine if mutations in *SLC26A11* impact symbiosis, we developed a method to electroporate two independent pairs of sgRNA/Cas9-complexes targeting *SLC26A11* to induce mutations in Aiptasia zygotes (Fig 5B). Successful mutation of *SLC26A11* was confirmed for each guide pair by genotyping larvae (fig. S10). Larvae were settled and metamorphosed into aposymbiotic polyps and inoculated with algal symbionts. After several weeks, we observed mosaic defects in symbiont density in mutant animals (11/21 guide pair 1; 4/33 guide pair 2) (Fig. 5C). The mosaic animals had regions with both low-symbiont density (light) and high-symbiont density (dark). Wild-type control polyps had uniform symbiosis patterns (15/15). We tested if mutations in *SLC26A11* caused the symbiosis defects by isolating gastrodermal tissue from dark and light tentacles from 12 mosaic animals and genotyping the sgRNA target sites in *SLC26A11*. The light tentacles harbored significantly more mutations than the dark tentacles for both guide pairs (Fig. 5D). We then performed whole-mount immunofluorescence staining of SLC26A11 in light and dark tentacles and found a significant decrease in SLC26A11 protein localization around symbiosomes in light tentacles (Fig. 5, E and F and fig. S11), indicating that loss of SLC26A11 protein around symbiosomes caused the symbiosis defects. Because inorganic carbon and sulfate supply is tightly associated with morphological and photosynthetic parameters in other free-living algae (*33*, *38–40*), we hypothesized that mutations in host *SLC26A11* would impact algal physiology. Indeed, we found that the algal symbionts in light tentacles had lower chlorophyll fluorescence intensity and larger algal cell size compared to those found in dark tentacles (fig. S12). Our results indicate that mutations in *SLC26A11* ablate its function on the symbiosome, leading to a reduction in symbiont density and defects in algal physiology likely caused by the decreased availability of inorganic carbon and sulfate to fuel photosynthesis and cell growth.

Photosymbiosis has evolved independently in Aiptasia and reef-building corals (*5*). To ask if SLC26A11 is also important for symbiosis in reef-building corals, we performed CRISPR/Cas9-based mutagenesis of the ortholog of *SLC26A11* in *G. fascicularis*. Over two laboratory spawning events (*41*), we injected 1-cell zygotes with Cas9-only or two independent pairs of sgRNA/Cas9 complexes targeting *SLC26A11* (Fig. 5G). We raised and settled the larvae into aposymbiotic juvenile polyps, which were then inoculated with algal symbionts. Strikingly, after two weeks, we found that the majority (13/28 guide pair 1; 21/22 guide pair 2) of sgRNA/Cas9-injected animals had significantly lower symbiont densities compared to both wild-type and Cas9-only-injected control polyps (Fig. 5H and fig. S13). We then performed high-throughput amplicon sequencing of PCR products spanning the sgRNA-target sites to quantify the mutation frequencies in each polyp. We found that high mutation rates correlated with lower symbiont density in sgRNA/Cas9-injected polyps (Fig. 5I). The algal symbionts in highly mutant polyps had lower algal chlorophyll fluorescence intensity and larger algal cell diameters for both guide pairs, mirroring our findings in Aiptasia (fig. S14). These results indicate that SLC26A11 has been repeatedly co-opted to play a critical role in symbiosis in two lineages that independently evolved photosymbiosis.

## Conclusion

We conclude that the symbiosome is a phagolysosome due to the abundance of lysosomal proteins, evidence of lysosomal fusion, and the negative impacts on symbiosis formation caused by the knockdown of key lysosomal machinery. In addition, our proteomics data revealed diverse digestive enzymes on the symbiosome suggesting phagolysosome-like digestive functions. However, the precise functions of these enzymes and how the algae persist in this seemingly hostile compartment remain unclear. The alga may inactivate the digestive machinery or may be generally resistant to digestion by the symbiosome. One possible explanation for the presence of digestive enzymes in the symbiosome comes from previous research that identified multiple layers of algal-derived membranes between the algae and the symbiosome membrane (*42*, *43*). These membranes are thought to be the result of algal ecdysis and contain lipids, carbohydrates, and proteins (*43*), which may be targeted for digestion by the symbiosome enzymes. The resulting products could be exported across the symbiosome through the diversity of transporters we identified in our proteome and may be an important source of photosynthate provided to the host. This mechanism may explain both the phagolysosomal identity of the symbiosome with robust digestive functions and the diversity of metabolites that are exchanged across the symbiosome.

The phagolysosomal identity of the symbiosome may explain how dinoflagellate algae (family Symbiodiniaceae) have repeatedly evolved photosymbioses across eukaryotic phyla (fig. S15A) (*4*, *44–48*). In the early evolution of these endosymbioses, the algae may have been phagocytosed from the environment and shuttled to phagolysosomes, consistent with the presence of acidified symbiosomes in several symbiotic species (fig. S15A). We hypothesize that an important step in the evolution of endosymbiosis formation is the survival of the algal symbiont in phagolysosome-like conditions. Once in phagolysosomes and not digested, the algae are in the nutrient centers of the host cells (*49*), providing an opportunity for the co-option of existing lysosomal machinery to export photosynthates and provide nutrients to the algae (fig. S15B). This co-option of lysosomal transporters for symbiosis may provide a relatively simple path for the repeated evolution of photosymbioses with dinoflagellate algae (fig. S15B).

Supporting this evolutionary model, we found that co-option of a lysosomal protein, SLC26A11, ensures the long-term maintenance of dinoflagellate algae in *G. fascicularis* and Aiptasia. Our finding that SLC26A11 is required for symbiosis suggests that host transport of sulfate and inorganic carbon is important for supporting the intracellular symbiont. SLC26A11 may fulfill a critical component of a hypothesized carbon concentration mechanism in the symbiosome. Previous research has implicated the lysosomal V-ATPase in symbiosome acidification, facilitating the conversion of bicarbonate into carbon dioxide in the symbiosome lumen. However, this proposed mechanism requires a bicarbonate transporter (*9*). As SLC26A11 is the only bicarbonate transporter detected on the symbiosome, we propose that SLC26A11 works synergistically with V-ATPase to concentrate inorganic carbon for intracellular algal photosynthesis (Fig. 5J). In this model, carbon concentration for photosymbiosis can be fulfilled by co-opting existing lysosomal proteins. Because SLC26A11 is required for symbiosis in two independently evolved endosymbiotic relationships, we provide evidence that the re-use of lysosomal machinery is a key step in the evolution of photosymbioses with dinoflagellate algae.

**Fig. 5.**
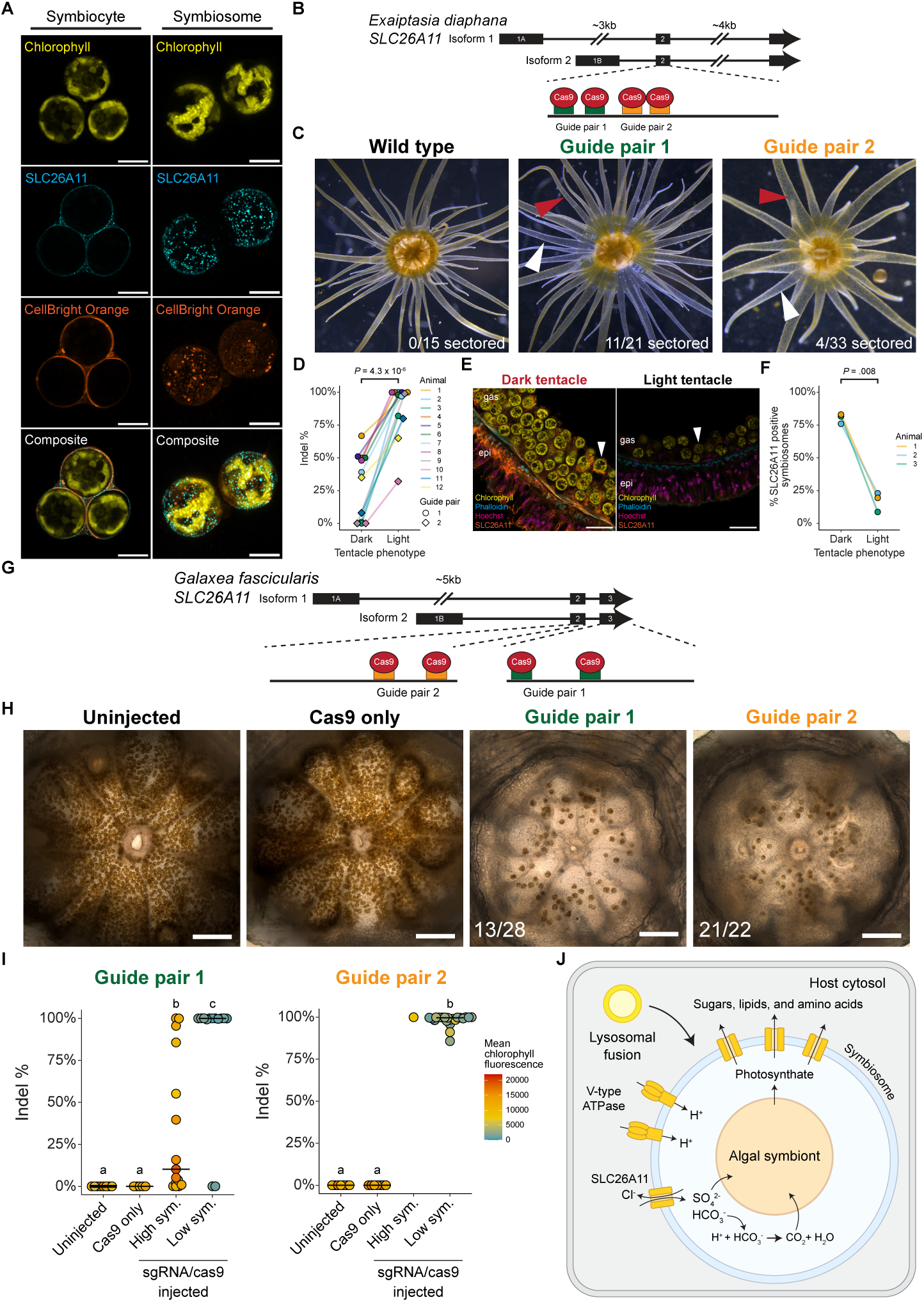
Mutations in *SLC26A11* causes symbiosis defects in Aiptasia and coral. **(A)** Immunofluorescence for SLC26A11 in Aiptasia symbiocytes and released symbiosomes. 5 μm scale bars. **(B)** Gene model for *SLC26A11* in Aiptasia with the locations of the two sgRNA target site pairs. **(C)** CRISPR/Cas9-mediated mutagenesis of *SLC26A11* results in mosaic Aiptasia with low-symbiont (light, white arrowheads) and high-symbiont (dark, red arrowheads) sectors. **(D)** *SLC26A11* mutation frequencies in gastrodermal tissue of paired light and dark tentacles from mosaic anemones. *P* values were determined by a paired t-test. **(E-F)** Immunofluorescence reveals reduced SLC26A11 around symbiosomes (white arrowheads) in light tentacles from animals treated with guide pair 1. 20 μm scale bars. *P* values determined by a paired t-test. **(G)** Gene model for *SLC26A11* in *Galaxea fascicularis* with the location of the two sgRNA target site pairs. **(H)** Animals injected with either pair of sgRNA/Cas9 complexes show reduced symbiosis compared to uninjected or Cas9-injected control animals. 100 μm scale bars. **(I)** Frequency of mutations and severity of symbiosis defects for each injection condition. Mean chlorophyll fluorescence is shown for each animal. Statistical significance was determined by a Kruskal-Wallis test followed by post-hoc Conover-Iman test (*P_adj_* < 0.05). **(J)** Model of SLC26A11’s role in symbiosis.

## Acknowledgements

We thank members of the Cleves lab for helpful comments on the manuscript. We thank the Carnegie Science operations team including Ted Cooper, Devance Reed, and David Ashwood. We thank the Carnegie Science Department of Embryology core facilities, including Mahmud Siddiqi, Ru-ching Hsia, Allison Pinder, Fred Tan, Xiaobin Zheng, and Joseph Tran for assistance and training in the execution of experiments. Finally, we thank the Carnegie Science Mass Spectrometry team, Tarabryn Grismer, Cao Son Trinh, Andres Reyes, and Shouling Xu for assistance in proteomics.

## Funding

Gordon and Betty Moore Foundation: GBMF12187 (PAC)

National Science Foundation: 2209203 (SM), 2128073 (PAC)

Revive and Restore Grant (PAC)

Pew Biomedical and Marine Fellow Award (PAC)

## Author contributions

Conceptualization: SM, PAC

Methodology: SM, CFH, NS, GPK, EMK, TE, PAC

Investigation: SM, CFH, NS, GPK, EMK, TE, PAC

Visualization: SM, CFH, NS, GPK, EMK, TE, PAC

Funding acquisition: SM, PAC

Project administration: PAC

Supervision: PAC

Writing – original draft: SM, PAC

Writing – review & editing: SM, CFH, NS, GPK, EKM, TE, PAC

## Competing interests

Authors declare they have no competing interests.

## Data and materials availability

Raw proteomics data is available in PRIDE repository XXX. Raw sequences and metadata have been deposited in the NCBI BioProject database (accession no. XXXXXX).

## Supplementary Materials

Materials and Methods

Figs. S1 to S16

Tables S1 to S15

References (50–191)

## Notes

### Competing Interest Statement

The authors have declared no competing interest.

