## Supplementary Materials for "Co-option of lysosomal machinery shapes the symbiosis supporting coral reefs"

**The PDF file includes:**

Materials and Methods
Figs. S1 to S16
Tables S1 to S16
References (50-191)

### Materials and Methods

*Animal husbandry and algal cultures*

Aiptasia (*sensu stricto Exaiptasia diaphana*) used in experiments were from the male clonal population CC7 symbiotic with the algal strain SSB01 (species *Breviolum minutum*) generated as previously described (*27*). A second clonal population of female Aiptasia, PLF3, was used only for spawning. Algal DNA was periodically sampled from CC7 animals and genotyped by sequencing PCR-amplified fragments of ITS2 as described previously to ensure that animals were not contaminated by other algal types (*27*). Unless otherwise stated, animals used in experiments were raised at 27 ºC, under a 12 h:12 h light:dark cycle under full spectrum Percival SciWhite LED lights (Percival, cat. no. AL-41L4) set to 25 μmol photons $m^{-2}$ $s^{-1}$, fed twice a week with freshly hatched *Artemia* nauplii in 1 L clear plastic containers filled with UV sterilized 5 μm filtered artificial seawater (FASW; Red Sea, cat. no. R11072). Water was changed with fresh FASW several hours after each feeding.

Nauplii and juveniles of *Tisbe* sp. copepods were fed to Aiptasia larvae to induce settlement (*50*). Starter cultures of *Tisbe* were obtained from Pod Your Reef (*Tisbe bimiensis* Reef Copepods) and Poseidon Reef Systems (Tisbe Copepod Starter Culture) and cultured continuously. The starter cultures were divided into four vessels with 8 L of FASW with continuous aeration through airstones. Each culture was fed 10 mL of RGCOMPLETE™ (Reef Nutrition) three times a week. The culture was harvested every 9 to 11 days by filtering the entire culture through 125 μm and 55 μm nylon mesh filters. The copepods that were captured by the 125 μm filter were returned to restart the culture with 8 L of FASW and 10 mL RGCOMPLETE™. The small copepods captured by the 55 μm mesh were further separated by size through 105 μm, 75 μm, and 55 μm nylon mesh filters. The 105 μm filter captured larger juvenile copepods that were discarded, the 75 μm filter captured larger nauplii and juvenile copepods, and the 55 μm filter captured small nauplii. Starting at 2 days post fertilization (dpf), Aiptasia larvae were first fed small nauplii five times a week during their first week of feeding, followed by larger nauplii and juvenile copepods five times a week during the second week. The larvae were then fed newly hatched *Artemia* nauplii five times a week until the larvae settled and metamorphosed into polyps (14-21 dpf). Larvae were given a complete water changes with fresh FASW in between each feeding.

SSB01 algal cultures, maintained axenic as previously described (*27*), were grown in IMK media at 27ºC, under a 12 h:12 h light:dark cycle under full-spectrum Percival SciWhite LED lights at 25 μmol photons $m^{-2}$ $s^{-1}$. Algal cultures were passaged to new media every four weeks. Cultures used in experiments were passaged two weeks prior to use and washed three times with FASW by pelleting for 1 min at 3000 x g before use. Algal concentration was determined using a GUAVA HT flow cytometer (Cytek) (*51*) and diluted with FASW as necessary.

*Generation of antibodies targeting Aiptasia proteins*

The RHBG antibody used to label symbiosomes has been described previously and was a kind gift from Dr. Manuel Aranda (*12*). Custom polyclonal antibodies were raised in rabbit by Genscript through their PolyExpress service against peptides for LAMP1A (AIPGENE27242), LAMP1B (AIPGENE27192), CTSB (AIPGENE22204), and truncated protein from the C-terminus of SLC26A11 (AIPGENE5335, AIPGENE5336). Detailed antigen target information is provided in table S6. Briefly, peptide or His-tagged protein targets were synthesized. For each antigen target, two rabbits were immunized with peptide and with keyhole limpet hemocyanin as a carrier protein or truncated SLC26A11 protein alone. Animals were sacrificed, sera collected and pooled, and antibodies were affinity-purified against their respective synthesized peptide or protein antigen targets.

To confirm the specificity of the custom antibodies, western blot analysis was performed on aposymbiotic and symbiotic Aiptasia polyp homogenate or aposymbiotic Aiptasia larvae homogenate (fig. S4). Briefly, three symbiotic or aposymbiotic polyps (1-2 mm oral disk diameter) or 330 aposymbiotic larvae were homogenized in 100 µL HU buffer (8M urea, 5% w/v SDS, 200 mM $NaHPO_4$ pH 6.8, 0.1 mM EDTA, 0.1% w/v bromophenol blue, 100 mM dithiothreitol, 1X cOmplete™ Mini EDTA-free Protease Inhibitor Cocktail; Roche, cat. no. 11836170001) for 2 min on ice using a motorized pestle (Kimble, cat. no. 749540-0000) and dounce (Kimble, cat. no. K749521-1500) in a 1.5 mL tube. Next, homogenized tissue was incubated at 95°C for 10 min to denature protein, followed by centrifugation at 14,000 x g for 10 min at 4°C to pellet cell debris. The supernatant was transferred to a new 1.5 mL tube, and 10 µL of the homogenate was directly loaded onto a stacking Tris-glycine SDS-PAGE gel (3.9% acrylamide stacking gel above an 8% acrylamide resolving gel). Proteins were separated by electrophoresis and transferred to a 0.2 µm PVDF membrane (Bio-Rad, cat. no. 1620177) by electrophoresis at 30 V overnight at 4°C in Towbin transfer buffer (25 mM Tris, 192 mM glycine, 20% v/v methanol).

PVDF membranes were stained with Ponceau S (Cell Signaling Technology, cat. no. 59803) for 10 min and washed five times with deionized water to confirm successful transfer of proteins. The stain was removed by replacing the water with 0.1M NaOH in deionized water and incubating for 2 min with gentle agitation on an orbital shaker. Next, the PVDF membrane was washed three times with deionized water, and the blot was blocked with a blocking buffer containing 5% Blotto non-fat dry milk (Santa Cruz Biotechnology, cat. no. NC9730946) in TBST (1% Tween-20 in TBS) for 2 hrs at room temperature (RT) with gentle agitation. The blocking buffer was removed and the membrane was incubated with a primary antibody solution containing 0.2 µg/mL of rabbit-α-LAMP1A, rabbit-α-LAMP1B, rabbit-α-CTSB, or rabbit-α-SLC26A11 antibody diluted in the blocking buffer overnight at 4°C with gentle agitation. Afterwards, the membrane was washed with three 15 min TBST washes at RT with gentle agitation before incubation with a secondary antibody solution containing a 1:10,000 dilution of horseradish peroxidase-linked goat-α-rabbit IgG antibody (Cell Signaling Technology, cat. no. 7074) in blocking buffer for 2 h at RT with gentle agitation. The solution was removed, and the membrane was washed with three 15 min TBST washes at RT with gentle agitation before dispensing 1 mL of SuperSignal™ West Femto Maximum Sensitivity Substrate (Thermo Fisher Scientific, cat. no. 34096) working solution directly onto the membrane. Blots were then imaged immediately on a LI-COR Odyssey Fc imager set to capture chemiluminescence signal.

The blots were then reprobed for alpha-tubulin as a protein loading control. Immediately after imaging, blots were rinsed briefly with TBST three times, then chemically stripped by incubating the blot in Restore™ Western Blot Stripping buffer (Thermo Fisher Scientific, cat. no. 21059) for 30 min at RT with gentle agitation. The strip buffer was removed, and the blot was washed with three 5 min TBS washes at RT with gentle agitation. Next, the blot was blocked with blocking buffer for 2 hrs at RT with gentle agitation. The blocking buffer was removed and the membrane was incubated with a primary antibody solution containing a 1:10,000 dilution of mouse-α-alpha-tubulin antibody (Cell Signaling Technology, cat. no. 3873) in blocking buffer overnight at 4°C with gentle agitation. Afterwards, the membrane was washed with three 15 min TBST washes at RT with gentle agitation before incubating with a secondary antibody solution containing a 1:10,000 dilution of horseradish peroxidase-linked goat-α-mouse IgG(H+L) antibody (Southern Biotech, cat. no. 1031-05) in blocking buffer for 2 h at RT with gentle agitation. The solution was removed, and the membrane was washed with three 15 min TBST washes before dispensing 1 mL of SuperSignal™ West Femto Maximum Sensitivity Substrate working solution directly onto the membrane. Next, blots were imaged immediately on a LI-COR Odyssey Fc imager set to capture chemiluminescence signal.

*Preparation of calcium-magnesium-free seawater*

To prepare calcium-magenesium free-seawater (CMFSW), a high concentration solution of CMFSW was first prepared containing 898 mM NaCl, 18 mM KCl, 66 mM $Na_2SO_4$, 4.3 mM $HCO_3$, 20 mM Tris-Cl pH 8 in Milli-Q® purified water. Next, high concentration CMFSW was diluted with Milli-Q® purified water until the solution measured 35 PPT on a digital refractometer (Milwaukee, cat. no. MA887), creating a working solution of CMFSW.

*Animal dissociation*

To dissociate Aiptasia polyps into single-cell suspensions, one to three whole anemones (pedal disk size ~10 mm) were incubated in 10 mL of dissociation buffer (4% w/v L-cysteine in CMFSW pH 8.5) with gentle agitation on an orbital shaker set to 85 RPM in a 60 x 15 mm plastic petri dish (Corning, cat. no. 351007). After approximately 1.5 h of incubation, the majority of epidermal tissue from the anemones dissolved away, leaving primarily gastrodermal tissue intact (fig. S16). Next, the remaining gastrodermal tissue was transferred with minimal buffer into a 5 mL conical tube, and fresh dissociation buffer was added to reach a total volume of 200 µL. Finally, the tissue was further dissociated into single cells by gently pipetting the sample 20 times through a P200 pipette set to 200 µL. The cells were then filtered through a 40 µm EASYstrainer cell strainer (Greiner Bio-One, cat no. 542140) to remove remaining tissue.

*Immunofluorescence and staining of live dissociated Aiptasia cells*

After dissociation, cells were diluted in an equal volume of a primary antibody solution containing 20 µg/mL of rabbit-α-RHBG antibody and 8% (w/v) BSA in CMFSW. Cells were incubated for 30 min at RT with gentle agitation, then centrifuged at 800 x g for 5 min to pellet cells. The supernatant was removed and cells were incubated with a secondary antibody solution containing 20 µg/mL of Goat α-Rabbit IgG (H+L) Highly Cross-Adsorbed Secondary Antibody, Alexa Fluor™ Plus 488 (Thermo Fisher Scientific, cat. no. A32731), 10 µM Calcein Violet (Thermo Fisher Scientific, cat. no. C34858), and 4% BSA in CMFSW. In some experiments, the membrane stain, 2 µg/mL of Nile Red (Sigma-Aldrich, cat. no. 19123), was also included into the secondary antibody solution for 30 min at RT with gentle agitation on an orbital shaker. Next, the cells were pelleted with an 800 x g centrifugation for 5 mins, the supernatant was discarded, and the cells were resuspended in CMFSW. Cells were mounted onto glass slides with #1.5 glass coverslips raised with clay feet or onto 35mm #1.5 glass bottom dishes (cat. no. D35-10-1.5-N; Cellvis, Mountain View, CA).

Stained cells were imaged with either an epifluorescence light microscope or a confocal microscope. Epifluorescence microscopy was performed on a Leica DM6 light microscope with 40X and 60X objectives under differential interference contrast (DIC) and fluorescence: DAPI (Ex BP 350/50; Em BP 460/50) for Calcein Violet, GFP (Ex BP 480/40; Em BP 527/30) for RHBG protein, and Cy5 (Ex BP 620/60; Em BP 700/75) for chlorophyll fluorescence. Super-resolution confocal microscopy was performed on Zeiss LSM 980 confocal with Airyscan 2 at 405 nm excitation (420-480 nm detection) for Calcein Violet, 488 nm excitation (495-578 nm detection) for RHBG protein, 560 nm excitation (574-620 nm detection) for Nile Red and 639 nm excitation (655-735 nm detection) for chlorophyll autofluorescence.

*Biochemical purification of the symbiosome*

Animals were sampled 2 h after the lights turned on during the 12 h:12 h light:dark cycle for symbiosome purification. Eighteen replicates, each consisting of five animals (pedal disk size ~10 mm), were incubated in 10 mL of dissociation buffer in 60 x 15 mm plastic petri dishes.

During incubation, 6 mL of dissociation buffer was replaced with 6 mL of fresh dissociation buffer every 30 min in each dish to ensure complete dissociation of the epidermal tissue. After 2 h of dissociation, removal of epidermal tissue was confirmed by microscopy. The resulting gastrodermal tissue from three dishes was transferred in minimal volume into a 5 mL tube for each biological replicate (n = 6). Next, ice-cold dissociation buffer was added to bring the total volume of each sample to 500 µL. Further dissociation of the gastrodermal tissue was performed by gently pipetting each sample 20 times through a P1000 pipette (Eppendorf, cat. no. 3121000120) set to 500 µL on ice. Next, 500 µL of ice-cold dissociation buffer was added to each sample. For each sample, 100 µL of the crude gastrodermal dissociation was aliquoted to be used as the control sample for proteomics, and the remaining 900 µL were further processed for symbiosome purification. While both control and symbiosome samples were processed in parallel, each sample preparation method will be described separately below.

For the control samples, 300 µL of ice-cold dissociation buffer was added and the samples were centrifuged at 100 × g for 5 min at 4°C. Next, the supernatant was removed, and the pellet was resuspended in 100 µL of ice-cold dissociation buffer. Afterwards, the samples were centrifuged at 100 × g for 5 min at 4°C, the supernatant was removed, and the pellet was resuspended in 200 µL of 0.4% NP-40 in 3.5X PBS. Then, each sample was passed through a 23-gauge needle (BD, cat. no. 305145) attached to a 3 mL syringe (BD, cat. no. 309657) three times to lyse the animal cells while keeping the algae intact. Two steps were performed to remove intact algae from the solution. First, the algae in each sample were pelleted by centrifuging at 2000 x g for 1 min at 4°C. Second, the resulting supernatant of each sample was loaded onto a 0.22 µm Ultrafree-MC centrifugal filter (Millipore, cat. no. UFC30GV0S) and centrifuged at 14,000 × g for 5 min at 4°C to further remove algal cells. The filtrate was collected, and 20 µL aliquots were taken from each sample for protein quantification. The remaining samples (~180 µL each) were then snap-frozen on dry ice.

For the symbiosome samples, gastrodermal cells were gently lysed by passing the sample three times through a 23-gauge needle attached to a 3 mL syringe. This step lysed the majority of the gastrodermal cells and caused the symbiosomes containing algae to be released from host cells. Next, 3 mL of ice-cold dissociation buffer was added to each sample. The symbiosomes containing algae were pelleted by centrifugation at 100 × g for 5 min at 4°C. The supernatant was removed, and the pellet was resuspended in 1 mL ice-cold dissociation buffer. Differential centrifugation was used to further purify released symbiosomes containing algae using a 40%-60%-100% discontinuous Percoll gradient. To prepare a 100% Percoll solution that was isotonic to 35 ppt seawater, 0.4 g of the dry components of CMFSW (26.24 g NaCl, 0.671 g KCl, and 4.687 g $Na_2SO_4$) was dissolved in 1 mL Milli-Q® purified water and added to 10.75 mL of Percoll (Sigma-Aldrich, cat. no. P4937). The 40% and 60% Percoll solutions were made by diluting the 100% isotonic Percoll solution with CMFSW. To make a 40%-60%-100% discontinuous gradient, 2 mL of each solution was added sequentially to a 15 mL conical tube. For each sample, the crude symbiosome was added to the top of the discontinuous gradient and centrifuged at 2000 x g for 5 min at 4°C. After centrifugation, host cell debris remained at the top of the 40% Percoll layer, while the released symbiosomes containing algae migrated to the 60%-100% Percoll boundary. To collect the released symbiosomes containing algae from this boundary, 4 mL of solution was first removed from the top of the gradient, then a P1000 with tip was directly inserted into the 60%-100% Percoll boundary, to extract 1 mL from this boundary. To confirm sample quality, 50 µL was aliquoted for staining with RHBG antibody and Calcein Violet as described above. Percoll was removed from the sample by adding 3 mL of CMFSW, centrifuging at 2000 x g for 1 min at 4°C to pellet, and discarding the supernatant. The resulting pellet was resuspended in 200 µL of 0.4% NP-40 in 3.5X PBS and passed through a 23-gauge needle three times to remove the symbiosome membrane from the algal cells. Next, the algae were removed from the sample, an aliquot was taken from the protein sample, and the protein sample flash frozen, following the same procedure described above for control samples.

An extra representative sample was prepared following the above procedure to quantify algal cell identity from aliquots taken at each major step of the protocol: (1) after cell dissociation, (2) after host cell lysis by passing the sample through a 23-gauge needle, (3) after Percoll density centrifugation, and (4) after stripping the symbiosome with 0.4% NP-40. Labeling was performed as described for immunofluorescence of dissociated cells and imaged on a Leica DM6 microscope at 40X magnification.

*Protein quantification*

We adopted an SDS-PAGE colloidal blue Coomassie staining method to measure the total protein mass of each sample. Samples and BSA standards were diluted with Laemmli buffer with 100 mM DTT, loaded onto a 10% acrylamide SDS-PAGE gel, and embedded into the gel by electrophoresis for15 min at 150 V. The gel was stained and fixed in colloidal blue (Invitrogen, cat. no. LC6025) following manufacturer's instructions. Gels were then visualized using an LI-COR Odyssey Fc imager using the 700 nm fluorescence detection channel. Sample protein concentrations were calculated by measuring the total lane fluorescence intensity at 700 nm and comparing it against standard curves generated from BSA standards.

*LC-MS/MS protein analysis*

A total of 4.5 µg of total protein from each control and symbiosome sample was diluted in Laemmli buffer with 100 mM DTT and loaded onto a 10% polyacrylamide Mini-Protean TGX gel (Bio-Rad, cat. no. 4561033). Samples were fixed and stained with colloidal blue following the manufacturer's instructions. Protein lanes were excised from the gel and frozen until further processing by in-gel tryptic digestion.

To perform in-gel digestion, gel samples were first washed several times with 25 mM ammonium bicarbonate/50% acetonitrile to equilibrate the gels. Next, the samples were reduced with 10 mM dithiothreitol, alkylated with 50 mM iodoacetamide, and dried using a speed vac. Trypsin solution was then added, and the samples were digested overnight at 37°C. The peptides were extracted by vortexing with 50% ACN/0.1% formic acid, then desalted using C18 ZipTips (Millipore, cat. no. ZTC18S).

LC-MS/MS was performed on an Orbitrap Eclipse mass spectrometer (Thermo Fisher Scientific) with an Easy LC 1200 UPLC liquid chromatography system (Thermo Fisher Scientific). Peptides were first trapped using a trapping column (Acclaim PepMap 100 C18 HPLC, 75 µm particle size, 2 cm bed length; Thermo Fisher Scientific, cat. no. 164946), then separated using an Aurora Ultimate™ 25×75 C18 UHPLC column (IonOpticks, cat. no. AUR3-25075C18). The flow rate was 300 nL/min. Peptides were eluted by a gradient from 3 to 28% solvent B (80% acetonitrile, 0.1% formic acid) over 106 min and from 28 to 44% solvent B over 15 min, followed by a 15 min wash at 90% solvent B.

Precursor scan was from mass-to-charge ratio (m/z) 375 to 1600 [resolution 120,000; automatic gain control (AGC) 200,000, maximum injection time 50 ms, Normalized AGC target 50%, radio frequency lens(%) 30], and the most intense multiply charged precursors were selected for fragmentation (resolution 15,000, AGC 5E4, maximum injection time 22 ms, isolation window 1.4 m/z, normalized AGC target 100%, include charge state = 2-8, cycle time 3 s). Peptides were fragmented using higher-energy collision dissociation with a normalized collision energy of 27. Dynamic exclusion was enabled for 30 s.

*Identification of differentially enriched proteins and prediction of protein function*

Raw spectral data were processed with Fragpipe (ver. 21.0) using "DDA+" as the data type and "LFQ-MBR" as the workflow settings and spectra were searched against the Aiptasia (ver. 1.0) (*52*) and *Breviolum minutum* (*53*) genomes. Including only the protein groups with Aiptasia Protein IDs, we normalized the spectral intensity of each protein group with Cyclic Loess normalization using the NormalyzerDE package (ver 1.18.1) in R with default settings. Only protein groups that were detected in at least three replicates from the control samples or least three replicates from the symbiosome samples were included in further analysis. Principal component analysis was conducted on a matrix of normalized spectral intensities using the *auto_plot* function of the ggfortify package (ver. 0.4.19) in R. Differentially enriched protein groups between control and symbiosome samples were identified using the NormalyzerDE package in R using default settings with an adjusted *P* value cutoff of 0.05 and $\log_2$ fold-change cutoff of 1. To simplify downstream analyses, the Protein ID of the protein group was used in downstream analysis to simplify data interpretation.

To quantify the proportion of proteins in the control- and symbiosome-enriched groups that are predicted to have at least one transmembrane domain, we analyzed every protein in the Aiptasia genome using Transmembrane Helices Hidden Markov Model (TMHMM) 2.0 under default parameters (*54*). To quantify the proportion of control- and symbiosome-enriched proteins that are transcriptionally upregulated in symbiotic compared to aposymbiotic adult anemones, we re-analyzed a previously published RNAseq dataset (*27*). Briefly, reads from symbiotic (strain CC7-SSB01) and aposymbiotic (strain CC7) animals at time 0 h from the time course experiment were aligned to the Aiptasia genome (version 1.0) using STAR (version 2.5.1b) under default alignment parameters. Read counts for each gene were generated using HTSeq (version 0.6.1) under default parameters. Raw read counts were then library-normalized using the function *counts* with the parameter *normalize = TRUE* in DESeq2. To identify differentially expressed genes, we used DESeq2 with default parameters to generate $\log_2$ fold-change expression ratios and adjusted *P* values. We then used this re-analyzed dataset to quantify the number of the control- and symbiosome-enriched proteins that were transcriptionally upregulated in symbiotic compared to aposymbiotic adult anemones (adjusted $P$ value $< 0.05$; $\log_2$ fold-change $> 1.0$). Next, using the number of transmembrane domain containing proteins and transcriptionally upregulated symbiosis genes genome-wide, we statistically determined enrichment of each in the control- and symbiosome-enriched proteins by performing pair-wise Chi-square tests using the *pairwiseNominalIndependence* function with default settings from rcompanion package (ver. 2.5.0) in R. Finally, we performed a cell-compartment gene-ontology (GO) enrichment analysis for control- and symbiosome-enriched proteins. To do this, we used the topGO package (ver. 2.60.1) in R and analyzed for enrichment of GO-terms from each control- and symbiosome-enriched protein list using the GO-term annotations from Baumgarten et. al. 2015 (*52*). Symbiosome-enriched protein function was predicted based on their function in other organisms (see table S1 for the full list of symbiosome-enriched proteins along with their predicted functions and corresponding citations).

*Immunofluorescence of live symbiosomes and fixed symbiocytes*

Symbiosomes containing algae were isolated and collected for immunofluorescence as described in the above procedure for symbiosome purification after Percoll density centrifugation. For staining, 50 µL of symbiosomes was extracted from the Percoll gradient and mixed with 50 µL of a primary antibody solution containing 20 µg/mL of a primary antibody (rabbit-α-LAMP1A, rabbit-α-LAMP1B, rabbit-α-CTSB, or rabbit-α-SLC26A11) and 8% BSA in CMFSW. Symbiosomes were incubated for 30 min at RT with gentle agitation on an orbital shaker, then centrifuged at 800 x g for 5 min to pellet symbiosomes. The supernatant was removed, and the symbiosomes were resuspended with a secondary antibody solution containing 20 µg/mL Goat α-Rabbit IgG (H+L) Highly Cross-Adsorbed Secondary Antibody, Alexa Fluor™ Plus 488 (Thermo Fisher Scientific, cat. no. A32731), 1:100 (v/v) CellBrite® Orange (Biotium, cat. no. 30022), and 4% BSA in CMFSW. Symbiosomes were washed twice with CMFSW by centrifugation at 800 x g for 5 min and resuspended in CMFSW. As controls, a no primary antibody solution containing only 8% BSA in CMFSW was used in the primary antibody step and cultured algae (strain SSB01) was also stained with each primary antibody.

Symbiocytes used for immunofluorescence were isolated using a modified version of the protocol used to isolate symbiosomes. First, 1 to 3 anemones were briefly washed twice in dissociation buffer, then incubated in 60 x 15 mm glass petri dishes (Corning, cat. no. 3160-60) with 10 mL dissociation buffer until epidermal tissue was separated from the gastrodermal tissue (approximately 30 min). The gastrodermal tissue was collected with minimal buffer into a 5 mL conical tube, and additional dissociation buffer was added to bring the volume up to 200 µL. The tissue was then further dissociated by repeatedly pipetting the sample through a P200 pipette ten times. Then, 800 µL of CMFSW was added to increase the sample volume and to dilute the dissociation buffer. The sample was then filtered through a 40 µm EASYstrainer cell strainer to remove remaining clumps of tissue. The filtrate was then loaded onto a 20%-40% iodixanol Optiprep gradient with 2 mL of each layer (Sigma-Aldrich, cat no. D1556) diluted in CMFSW. To prepare the gradient, the stock Optiprep solution (60% iodixanol in water) was diluted to 20% and 40% iodixanol with CMFSW. To make a 20%-40% Optiprep gradient, 2 mL of each solution was added sequentially to a 5 mL conical tube. The gradient loaded with the filtrate was then centrifuged at 1000 x g for 1 min. Non-symbiotic host cells remained at the top of the gradient above the 20% layer, and symbiocytes and algae were collected from the 20%-40% boundary.

The recovered symbiocytes in Optiprep solution were diluted in an equal volume of FASW to allow the cells to begin to sink, and 100 µL of the cells were quickly transferred to a 35mm #1.5 glass bottom dish (Cellvis, cat no. D35-10-1.5-N) coated with Gibco poly-D-lysine (Thermo Fisher Scientific, cat no. A3890401) following the manufacturer's instructions. Cells were allowed to settle and adhere onto the glass dish for 10 min at RT. Next, the solution was removed and replaced with 1% paraformaldehyde and 0.005% glutaraldehyde in FASW, and cells were allowed to fix for 10 min at RT with gentle agitation on an orbital shaker. The fixed and adhered cells were then washed with three 5 min FASW washes with gentle agitation at RT. For the α-CTSB antibody stain and its respective controls, cells were then incubated in an antigen-retrieval buffer containing 10 mM citrate and 0.1% saponin in 1X PBS for 10 min at 95°C. The cells that underwent antigen retrieval were then washed with three 5 min 1X PBS washes at RT with gentle agitation. For all antibody stains, cells were then incubated with a permeabilization/blocking buffer containing 1% BSA and 0.1% saponin in 1X PBS for 1 h at RT with gentle agitation. The buffer was removed, then replaced with a primary antibody solution containing an antibody in permeabilization/blocking buffer and incubated overnight at 4°C with gentle agitation. Antibody final concentrations used were 10 µg/mL for rabbit-α-LAMP1A, rabbit-α-LAMP1B, rabbit-α-CTSB, rabbit-α-IgG isotype control (Genscript, cat. no. A01008), and 0.685 µg/mL for rabbit-α-SLC26A11. Cells were then washed with three 10 min permeabilization/blocking buffer washes at RT with gentle agitation. The buffer was then replaced with a secondary antibody solution containing 4 µg/mL Goat α-Rabbit IgG (H+L) Highly Cross-Adsorbed Secondary Antibody Alexa Fluor™ Plus 488, 1:100 (v/v) CellBrite® Orange to stain membranes and 10 µg/mL Hoechst 33342 to stain DNA (Thermo Fisher Scientific, cat. no. 62249) and incubated for 2 h at RT in the dark with gentle agitation. Cells were then washed with three 10 min permeabilization/blocking buffer washes, followed by two 10 min 1X PBS washes. Finally, the cells were mounted in 1X PBS, a coverslip was placed over the well and imaged immediately.

Live symbiosomes and fixed symbiocytes were imaged on a Zeiss LSM 980 confocal with Airyscan 2 at 405 nm excitation (420-480 nm detection) for Hoechst 33342 (when used), 488 nm excitation (495-578 nm detection) for antibody detection, 560 nm excitation (574-620 nm detection) for CellBrite® Orange, and 639 nm excitation (655-735 nm detection) for chlorophyll autofluorescence. Vesicle diameter was manually quantified using FIJI (ver. 2.14.0) by measuring the distance across the widest part of each vesicle from 17-53 vesicles in each sample.

Live symbiosome and cultured algae used as controls (fig. S5) were imaged on a Leica DM6 light microscope with 40X objectives under differential interference contrast (DIC) and fluorescence: GFP (Ex BP 480/40; Em BP 527/30) for antibody, and Cy5 (Ex BP 620/60; Em BP 700/75) for chlorophyll fluorescence.

*Lysotracker stain*

Small anemones (approximately 1-3 mm pedal disc diameter) were placed into 2 mL round-bottom tubes with 1 mL of staining solution containing 100 nM Lysotracker Green in FASW. Animals were stained for 30 min at RT in the dark with gentle agitation. Animals were then directly mounted onto glass slides with #1.5 coverslips raised with clay feet and to allow for clearer visualization of symbiocytes, the tissue was gently compressed by the coverslips immediately before imaging. Mounted animals were imaged on a Zeiss LSM 980 confocal with Airyscan 2 at 488 nm excitation (495-578 nm detection) for Lysotracker Green and 639 nm excitation (655-735 nm detection) for chlorophyll autofluorescence.

*Design and generation of short-hairpin RNAs*

Short hairpin RNAs (shRNAs) targeting genes of interest were designed and synthesized using previously described methods (*30*). Hairpins were designed using siRNA Wizard software v3.1. (https://www.invivogen.com/sirnawizard/) against target mRNA sequences with the following parameters: 19-bp target motifs starting with one or two guanines and between 40-50% overall GC content. Designed shRNAs were checked for potential off-targets using Bowtie version 1.3.0 (*55*) against the Aiptasia genome and gene models (*52*), and sequences that had off-targets with three or fewer mismatches were excluded. The shRNA sequence was designed as follows: the T7 promoter sequence, followed by the sense target sequence, a looping linker sequence, the antisense sequence, and two terminal thymidines (table S7). Two shRNAs were designed per gene target to be co-electroporated.

To synthesize each shRNA, two single-stranded DNA templates, one for the designed shRNA sequence, and another of its reverse complement, were ordered as oligonucleotides through Integrated DNA Technologies. The oligonucleotides were reconstituted to 100 µM in duplexing buffer (30 mM HEPES pH 7.5, 100 µM potassium acetate). Before *in vitro* transcription, the two DNA oligos were annealed by combining 5 µL of each oligonucleotide followed by denaturation for 5 min at 98°C, then immediately cooled to 24°C for 10 min. For *in vitro* transcription, 2 µL of the annealed template was used in a 60 µL *in vitro* transcription reaction with a HiScribe T7 Quick High Yield RNA Synthesis kit (New England Biolabs, cat. no. E2040L). After an overnight incubation, the RNA samples were diluted with 60 µL of nuclease-free water, and 4 µL of DNAse-I (New England Biolabs, cat. no. M0303L) was added to each reaction and incubated at 37°C for 30 min. After the incubation, the resulting RNA was purified using an RNA Clean and Concentrator-25 kit (Zymo Research, cat. no. R1018) following manufacturer's instructions. RNA size and quality was determined on a 2% agarose gel, and RNA concentration was determined using a Nanodrop spectrophotometer. RNA was stored at -80°C. Scrambled control shRNAs were created for each guide pair sharing similar GC content that do not bind to Aiptasia genes in the genome or gene models as determined by Bowtie and synthesized as described above (table S8).

*Short-hairpin RNAi knockdown experiments in* Aiptasia

Animals were spawned (*56*) (dx.doi.org/10.17504/protocols.io.ru2d6ye), and zygotes were electroporated to deliver shRNA using previously described methods (*30*). Briefly, eggs were fertilized within 30 min after spawning and collected into glass bowls (Carolina biological, cat. no. 741000). Approximately 30 µL of zygotes (containing 300-1000 individuals) were then mixed with 50 µL of 30% w/v Ficoll in FASW and 20 µL of shRNA containing 30 µg of each hairpin. The solution was mixed and transferred to a Gene Pulser/Micropulser Electroporation Cuvette (Bio-Rad, cat. no. 1652088) and was immediately electroporated using a NEPA 21 Type II electroporator (Nepa Gene) using the following conditions: poring pulse at 80.0V, 25 ms pulse length, 99.0 ms pulse interval, 5 pulses, and 40% decay rate with no transfer pulse. Electroporated zygotes were immediately recovered by adding 2 mL of FASW into the cuvette, then transferred to 6-well plates containing 5 mL FASW. A complete water change was given to the larvae 24 h post-fertilization.

To verify knockdown efficiency, larvae were sampled for RNA extraction 2 dpf. Exactly 220 larvae were transferred into a 1.5 mL tube with minimal seawater and placed on ice for 2 min to halt swimming. The tube was then briefly spun down to gather the larvae to the bottom of the tube. Next, the seawater was removed and replaced with 1 mL of TRIzol (Thermo Fisher Scientific, cat. no. 15596026). Afterwards, 100 µL of 0.5 mm diameter glass beads (Biospec, cat. no.11079105) were added and the sample was vortexed for 30 s. Then, 100 µL of bromochloropropane was added and the sample was vortexed again for 30 s, quickly spun down, and incubated for 10 min at RT. The sample was then centrifuged at 16,000 x g for 15 min at 4°C. The upper aqueous layer (~400 µL) containing RNA was transferred to a new tube. Next, 200 µL of 100% isopropanol and 200 µL of High-Salt solution for Precipitation (Plant) (Takara Bio, cat. no. 9193) was added to the RNA sample, vortexed briefly, then incubated at -20°C for 48 h to precipitate RNA. Afterwards, the sample was centrifuged at 16,000 x g for 15 min at 4°C and the supernatant discarded. Next, the pelleted RNA was washed with 1 mL of ice-cold 75% ethanol, centrifuged at 7,600 x g for 5 min at 4°C and the supernatant discarded. The ethanol wash step was then repeated once more. The RNA pellet was air dried to remove excess ethanol, and the RNA was dissolved in 50 µL of nuclease-free water. The sample was further purified using a Quick-RNA Microprep Plus Kit (Zymo Research, cat no. R1050) following the manufacturer's instructions including the on-column DNase I step. The RNA was eluted in 15 µL of nuclease-free water, quantified using a Nanodrop Spectrophotometer, and stored at −80°C. RNA samples were diluted to the lowest concentration sample with nuclease-free water, and an identical amount of RNA (10-20 ng) was used for cDNA synthesis using an iScript cDNA kit (Bio-Rad, cat. no. 1708891) following the manufacturer's instructions. Briefly, samples were incubated for 5 min at 25°C, 20 min at 46°C, and 1 min at 95°C. The resulting cDNA was diluted 1:10 in nuclease-free water to make a stock solution stored at -20°C. RT-qPCR was then performed using the primers described in table S9. Three or four technical replicate qPCRs were performed for each biological replicate. To prepare the PCR reactions, 2 µL of diluted cDNA was added to a mix of 5 µL of SsoAdvanced Universal SYBR Green Supermix (Bio-Rad, cat. no. 1725270), 0.375 µL of each primer, and 2.25 µL nuclease-free water in a well of a white PCR plate (Brandtech, cat. no. 781367). PCR was performed on a CFX96 Real-Time System (Bio-Rad) using the following cycling parameters: 30 s at 95°C; 45 cycles of 10 s at 95°C, and 45 s at 58°C; and a final melting step of 60 cycles of 5 s with a temperature ramp from 65°C to 95°C in 0.5°C increments per cycle.

In addition to the PCRs for the target genes, parallel PCRs of a housekeeping gene *RPS7* (AIPGENE28828) (*57*), were performed on each sample to be used for normalization and the calculation of relative expression levels of the target genes. To determine relative expression levels of target genes, threshold cycles (CT) for each sample were first internally normalized to that of the housekeeping gene using the following equation: $\Delta CT = CT_{\text{gene of interest}} - CT_{RPS7}$. Fold-changes in expression levels were then calculated using the $2^{-\Delta\Delta CT}$ method (*58*) relative to the appropriate controls.

To inoculate larvae with algae, approximately 100 larvae (2 dpf) were transferred to each of three replicate 60 x 15 mm glass dishes containing 10 mL of strain SSB01 algae at 150,000 cells/mL for each condition in FASW. Larvae were incubated for 3 d at 27°C on a 12 h:12 h light:dark cycle. Afterwards, the larvae were transferred to a new glass dish with 10 mL FASW and incubated for 3 d without algae at 27°C on a 12 h:12 h light:dark cycle. After 3 d, larvae were transferred to a 4-well plate (Thermo Fisher Scientific, cat no. 179820) in ~40 µL total volume and fixed by adding 40 µL fixative (4% paraformaldehyde and 0.02% glutaraldehyde in FASW) for 5 min at RT with gentle agitation on an orbital shaker. The solution was removed and replaced with 40 µL of fresh fixative and incubated for another 5 min at RT with gentle agitation. Next, the solution was removed and replaced with 80 µL of 4% paraformaldehyde in FASW and incubated for 1 h at 4°C. The larvae were then washed with five 5 min washes using 500 µL ice-cold PBSTw (0.1% Tween-20 in 1X PBS) with gentle agitation. Fixed larvae were then stained with a 300 µL solution containing 1:400 Alexa Fluor™ 488 Phalloidin (Thermo Fisher Scientific, cat. no. A12379) for 1 h at RT with gentle agitation. Stained larvae were then washed with two 15 min washes using 300 µL ice-cold PBSTw at RT with gentle agitation. Samples were then mounted in 70% glycerol in 10X PBS onto a glass slide with SecureSeal™ Imaging Spacers (Grace Bio-Labs, cat. no. SS10X6.35) and with #1.5 glass coverslips.

To quantify the proportion of larvae that had at least one symbiont within their gastrodermal tissue, the samples were imaged and manually scored under fluorescence microscopy using a Leica DM6 light microscope with a 10X or 20X objective. Z-stack images were taken under GFP (Ex BP 480/40; Em BP 527/30) for Phalloidin and Cy5 (Ex BP 620/60; Em BP 700/75) for chlorophyll fluorescence.

*Design and generation of single-guide RNAs targeting SLC26A11*

The Aiptasia transporter, *SLC26A11*, was identified as symbiosome-enriched from the proteomics experiments. To confirm the gene's orthology to human *SLC26A11*, we performed a reciprocal BLASTp search by first performing a search against the human proteome using default settings with AIPGENE5335 as the query, resulting in the top search hit being human SLC26A11 (NP_001159819.1). We then performed a second BLASTp search against the Aiptasia proteome using the human SLC26A11 as the query, resulting in the top search hit being the Aiptasia SLC26A11. Aiptasia *SLC26A11* has two predicted splice isoforms (AIPGENE5335 and AIPGENE5336) in the genome. Each isoform has three predicted exons, with the first exon differing between the two isoforms. To verify the gene model for *SLC26A11*, we predicted the number of transmembrane domains for the gene using DeepTMHMM (ver. 1.0.44) (*59*) and compared it to the prediction for *SLC26A11* in humans (Uniprot accession: Q86WA9). Each SLC26A11 isoform in Aiptasia had the expected 12 transmembrane domains consistent with the human SLC26A11 protein. To identify *SLC26A11* orthologs in *G. fascicularis*, we performed a BLASTp search against the *G. fascicularis* genome (*60*, *61*) using the AIPGENE5335 protein sequence as a query using default settings. This search identified four matches in the *G. fascicularis* proteome (gfas1.m1.6060.m1, gfas1.m1.3149.m1,, gfas1.m1.3154.m1, gfas1.m1.3153.m1), where the top hit was gfas1.m1.6060.m1 (table S10). A BLASTp search using the gfas1.m1.6060.m1 protein sequence against the Aiptasia genome protein database identified AIPGENE5336 as the reciprocal best BLAST match, supporting one-to-one orthology of these two proteins.

The *G. fascicularis SLC26A11* gene model predicted only one isoform with four exons. DeepTMHMM analysis of the protein predicted 16 transmembrane domains rather than the expected 12. We split exons 1 and 2 of this gene model to produce two isoforms that each shared exons 3 and 4, mirroring the Aiptasia *SLC26A11* gene model. DeepTMHMM predicts 12 transmembrane domains for each isoform, supporting these new gene models (table S10). In both Aiptasia and *G. fascicularis*, we designed two pairs of single-guide RNAs (sgRNAs) targeting the last two exons, which are shared in both isoforms of *SLC26A11* in each species.

To design sgRNAs for CRISPR/Cas9 mutagenesis, DNA sequences of exon 2 for Aiptasia and the last two exons for *G. fascicularis* were processed through the sgRNA target site prediction software, CHOPCHOP (ver. 3.0) (*62*), using default parameters. The list of predicted sgRNA target sites for each exon was then aligned to its respective genome and checked for predicted off-target binding in the genome using the following steps. First, the genome index was generated using the command *bowtie_build* from Rbowtie (ver. 1.48.0) package in R. Second, sgRNA sites were then aligned to their respective genomes using the command *runCrisprBowtie* from crisprBowtie (ver. 1.12.0) package in R with the following parameters: crisprNuclease = spCas9, n_mismatch = 3, and canonical = TRUE). Single-guide RNA sites with predicted off-target alignments with three or fewer mismatches were discarded. We then selected two pairs of sgRNAs for each species that ranked high on the efficiency score and ordered synthesized sgRNAs with 'standard modifications' from Synthego. Sequences of the sgRNA-target sites are provided in table S12.

*CRISPR/Cas9 mutagenesis experiments in Aiptasia*

Single-guide RNAs from Synthego were resuspended in nuclease-free water to a final concentration of 2 µg/µL. To make CRISPR/Cas9 ribonucleoprotein (RNP) complexes for electroporation, each sgRNA was combined with Cas9 protein by mixing 8 µL of resuspended sgRNA, 2 µL of Alt-R™ S.p. Cas9 Nuclease at 10 µg/µL (Integrated DNA Technologies, cat. no. 1081059), and 1.1 µL of 10X complexing buffer (200 mM Hepes, pH 7.3, 1.5 M KCl), which was then incubated at 37°C for 15 minutes. Next, 10 µL of each RNP complex for each sgRNA pair were mixed together, along with 2.5 µL of non-homologous 400 µM single-stranded oligodeoxynucleotide (ssODN, TGGTTACAACTTCTTTATTGCTGCCCTTGTTGACTAGG TAATCACTCGAGTATCAGCCTGCCAGTGGGCTACCATCTTGTCCGGTGT). The non-homologous ssODN was added to increase the delivery of RNP complexes into cells during electroporation (*63*).

To make the electroporation mixture, 30 µL of zygotes (containing 300-1000 individuals) were mixed with 50 µL of 30% w/v Ficoll in FASW and 20 µL of the paired RNP complexes including ssODN. The zygote solution was transferred into a cuvette, electroporated, and the transformed zygotes were washed as described above for delivery of shRNAs. After 24 h, the larvae were transferred to a new dish of FASW. At 2 dpf, DNA was extracted from eight larvae to assess mutation frequency around each guide site (DNA extraction methods are described in the next section). Starting at 2 dpf, the larvae were fed *Tisbe* nauplii as described above to induce settlement and metamorphosis. After ~90% of larvae developed into polyps (~14-21 days), animals were inoculated with cultured SSB01 algae at 10,000 cells/mL concentration. After three days of algae exposure, the animals were rinsed by replacing the FASW with fresh FASW and any remaining larvae were allowed to complete metamorphosis. The animals were maintained in standard culture conditions (see above) and fed freshly hatched Artemia nauplii twice a week with water changes performed after each feeding. The polyps were screened daily for symbiosis defects under a Leica Ivesta 3 Stereomicroscope. Between 14-21 days post-inoculation (dpi), polyps began to display mosaic differences in symbiont densities, which persisted over time. At 22 dpi (for guide pair 1) and 62 dpi (for guide pair 2), polyps were scored for mosaic differences in symbiont density with both white light illumination and by fluorescence microscopy using a GFP long-pass filter (400-455 nm excitation, 480 long-pass emission filter) using a Leica M165 FC stereoscope. Animals were considered to exhibit a mosaic phenotype if they had inconsistencies in symbiont densities in a radial pattern that started from the mouth and extended to the tentacles.

*Genotyping SLC26A11 mutation frequency in Aiptasia*

Individual polyps with mosaic phenotypes were transferred to separate 50 Dram Clear Polystyrene Plastic Vials (Thornton Plastics, cat. no. 50U) containing 140 mL of FASW. To determine if mutations in *SLC26A11* correlated with the mosaic differences in symbiont density, we quantified mutation frequencies in the gastroderm of tentacles with high-symbiont density (“dark tentacle”) and low-symbiont density (“light tentacle”) from individual mosaic polyps. Using microdissection scissors (Excelta, cat. no. 349B), we removed one dark and one light tentacle from 8 animals for guide pair 1 and 4 animals for guide pair 2. Each tentacle clip was placed in a small petri dish with 3 mL of dissociation buffer (4% L-cysteine in CMFSW, pH 8.5) and incubated at RT with gentle shaking for one hour to remove the epidermal cell layer.

To extract DNA from larvae 2 dpf after electroporation with sgRNA and from the resulting gastrodermal tissue of each tentacle, individual larvae or tentacles were transferred in a 1 µL volume into a 200 µL tube containing 9 µL of lysis buffer (10 mM Tris pH 8.3, 50 mM KCl, 50 mM $MgCl_2$ 0.3% Tween-20, and 0.3% NP-40). The samples were then incubated at 94°C for 20 min. Next, 0.5 µL of 20 mg/mL proteinase K (New England Biolabs, cat. no. P8107S) was added and incubated at 55°C to digest proteins. Afterwards, the samples were incubated at 94°C for 20 min to deactivate the enzyme. This crude DNA extract was then used directly for PCR as template.

PCR primers (Forward: AATACCCGCTACCAATCCCG; Reverse: CTGCTTGAAAAA TGTCGGTCACA) were designed to amplify a 564 bp product centered around all the four sgRNA target sites. To amplify this region, 1 µL of crude DNA extract was added to a PCR mix containing 12.5 µL of 2X GoTaq® Green Master Mix, 1 µL of 10 µM forward primer, 1 µL of 10 µM reverse primer, and 10.5 µL of nuclease-free water. The PCR reaction conditions were as follows: one cycle of 95°C for 2 min; 30 cycles of 95°C for 20 s, 59°C for 30 s, and 70°C for 10 s; one cycle of 70°C for 10 min. The size and quality of the resulting PCR product was confirmed with electrophoresis on a 1% agarose gel. Unpurified PCR products were purified and Sanger sequenced using GENEWIZ’s Sanger sequencing service for unpurified PCR products. Mutation frequences were determined for each PCR product by deconvoluting Sanger traces using Synthego’s Inference of CRISPR Edits software (https://ice.editco.bio/).

*Whole-mount immunofluorescence of Aiptasia tentacles*

To localize SLC26A11 protein, we performed whole-mount immunofluorescence in tentacles of Aiptasia with mosaic phenotypes. One light- and one dark-colored tentacle was removed from each of three mosaic animals (from sgRNA pair 1), then fixed in a well of a 4-well plate with 500 µL of 4% paraformaldehyde in FASW, and incubated for 30 min at RT with gentle agitation on an orbital shaker. After fixation, the tentacles were washed with three 5 min washes using 500 µL of PBST (0.2% Triton X-100 in 1X PBS). Next, fixed tentacles were manually cut into round transverse sections with a scalpel under a Leica Ivesta 3 Stereomicroscope. Tentacle sections were placed in 500 µL of permeabilization/block buffer containing 1% DMSO and 10% normal goat serum in PBST and incubated at RT for 2 h with gentle agitation. The buffer was then removed and replaced with 500 µL of primary antibody solution containing 0.685 µg/mL of rabbit-α-SLC26A11 antibody in permeabilization/block buffer and incubated at 4°C overnight with gentle agitation. The antibody buffer was removed and briefly washed twice with 500 µL of PBSTw, followed by three additional 20 min washes using 500 µL of PBSTw at RT with gentle agitation. The buffer was removed and replaced with 500 µL of secondary antibody solution containing 20 µg/mL Goat αRabbit IgG (H+L) Highly Cross-Adsorbed Secondary Antibody, Alexa Fluor™ Plus 555 (Thermo Fisher Scientific, cat. no. A32732), 1:200 Alexa Fluor™ 488 Phalloidin, 10 µg/mL Hoechst 33342, and 10% normal goat serum in PBST, and incubated for 2 h at RT with gentle agitation. The buffer was removed, and samples briefly washed twice with 500 µL of PBSTw, followed by three additional 20 min washes using 500 µL of PBSTw at RT with gentle agitation. The wash buffer was removed, and samples were incubated in 15 µL of Ce3D™ clearing solution (BioLegend, cat. no. 422703) for 5 min at RT. Afterwards, the samples were transferred to a glass slide in 1 µL volume and mounted with 10 µL of 80% glycerol in 1X PBS. Next, the sample was covered with a #1.5 coverslip raised with clay feet on each corner of the coverslip to minimize compression.

Prepared samples were immediately imaged on a Zeiss LSM 980 confocal with Airyscan 2 at 405 nm excitation (420-480 nm detection) for Hoechst 33342, 488 nm excitation (495-578 nm detection) for Phalloidin, 560 nm excitation (574-620 nm detection) for SLC26A11 antibody, and 639 nm excitation (655-735 nm detection) for chlorophyll autofluorescence. To quantify SLC26A11 abundance, we used FIJI to manually count the proportion of symbiosomes that had a positive signal for SLC26A11 antibody from 3-5 sections for each tentacle.

*CRISPR/Cas9 mutagenesis experiments in* Galaxea fascicularis

Complexing of sgRNA to Cas9 was performed as described for Aiptasia but with slight modifications. First, the Alt-R™ S.p. Cas9 Nuclease (10 µg/uL) was diluted with complexing buffer to 3 µg/µL. Then, 1.5 µL of sgRNA, diluted to 2 µg/µL with nuclease-free water, was mixed with 1.5 uL of diluted Cas9 enzyme in 3 µL total volume and incubated at 37°C for 15 min to complex. To combine pairs of sgRNA/Cas9 complex, 3 uL of each sgRNA/Cas9 was then mixed together. As a control, a Cas9 protein only solution was prepared by mixing 3 µL of diluted Cas9 enzyme and 3 µL of nuclease-free water and incubating at 37°C for 15 min.

*Galaxea fascicularis* collected from the Great Barrier Reef were induced to spawn in February 2025 in laboratory aquaria with programmable temperature and light controls as described in Swinhoe et al. 2025. Synchronous spawning of 6–11 coral colonies occurred on February 26 and 27. Gamete collection, washing, and crosses were performed as previously described in Swinhoe et al, 2025. Briefly, during each of these spawning nights, gametes from individual parental colonies were collected, rinsed, and isolated in plastic bowls at 27°C until fertilizations were performed. Staggered crosses of a mixture of all eggs and all sperm were performed for microinjection each hour for five hours.

Microinjection was used to deliver sgRNA/Cas9-complexes and Cas9-only control reagents to *G. fascicularis* zygotes as previously described (*41*) except for the use of a blue dye as an injection indicator that allows for sorting of successfully injected zygotes at 1 h post injection under white light. Injection solutions were prepared as follows: 5 µL of paired sgRNA/Cas9 or Cas9-only solution, 2.5 µL of complexing buffer, and 0.5 µL of 0.4 µm sterile-filtered FD&C Blue #1 (40.76 mg/mL in water; Spectrum Chemical, cat. no. FD110). Guide pair 1 and Cas9-only control were injected on February 26th and Guide pair 2 and Cas9-only control were injected on February 27th. Each day, approximately 400-500 1-cell zygotes were injected with sgRNA/Cas9 complexes or Cas9-only solutions. Uninjected 1-cell zygotes from the same fertilizations were collected as control animals. Immediately after injection, larvae were sorted as successfully injected by the presence of the blue injection indicator. Larvae were housed in plastic deli containers and incubated in standard culture conditions described above. Details of injections (number of zygotes injected, survival at 24 hpf, blue vs non-blue animals) are available in table S13.

Successfully injected larvae (sgRNA/Cas9 and Cas9 Only) and uninjected control larvae were induced to settle starting 1 dpf. For each class, groups of 10 - 30 animals were placed in a 500 µL or 1 mL drop of FASW centered in 60 x 15 mm petri dishes to prevent larvae from settling on the corners of the dish (see table S14 for details). The dishes were covered with lids, placed in a humidity chamber at RT. Three times a week, half of the water was replaced with fresh FASW. By 23- and 22-days post fertilization (for spawn 1 and 2, respectively), ~90% of the larvae had settled, and the dishes were transferred to a plastic tub containing 3 L of FASW, which was inoculated with 10,000 cells/mL of SSB01 culture. At 7 dpi and 14 dpi, the dishes were rinsed with FASW, and the water in the tub replaced with fresh FASW.

At 14 dpi, after washes, animals were imaged on a Leica DM6 light microscope using a 16X water immersion objective that was immersed directly into the petri dish containing seawater. Z-stack images were taken under DIC, GFP (Ex BP 480/40; Em BP 527/30) for coral autofluorescence and Cy5 (Ex BP 620/60; Em BP 700/75) for chlorophyll fluorescence. FIJI software was used to generate maximum intensity projections of each z-stack image using the *Z project* function. Images were scored blindly and independently by three people (authors S.M., C.F.H., and G.P.K.) for symbiont density determined from the chlorophyll channel as “High” or “Low”. The symbiont density scores strictly ignored chlorophyll intensity and was instead determined by the density of symbionts. Next, mean chlorophyll fluorescence in each juvenile coral was quantified by manually selecting the coral oral disk and tentacles in the DIC channel and measuring the fluorescence intensity in the Cy5 channel (fig. S13). Animals with developmental deformities were discarded from subsequent analysis. Each juvenile animal was tracked using a map of the petri dish for data analyses and interpretation.

*Genotyping SLC26A11 mutation frequency in* Galaxea fascicularis

Animals were sacrificed 18 dpi by scraping individual animals off of the petri dish with a P1000 pipette. Animals were transferred into individual 1.5 mL tubes and the seawater was removed. The samples were immediately washed once with 500 µL of 100% ethanol. The solution was removed, then replaced with 500 µL of fresh 100% ethanol and stored at -20°C. DNA extractions were performed as previously described (*64*). Briefly, ethanol was removed from each sample, replaced with 750 µL of lysis buffer (100 mM Tris pH 9.0, 100 mM NaCl, 100 mM EDTA, 1% SDS) and 20 µL of Proteinase K (20 mg/mL), and incubated at 65°C for 2 h. Next, 187.5 µL of 5 M KOAc was added to each sample, vortexed, and put on ice for 10 min. The samples were centrifuged at 16,100 x g for 20 min at RT. Next, the supernatant was transferred to a new tube and the pellet discarded. Afterwards, 10 µL of glycogen and 700 µL of 100% isopropanol were added and incubated for 30 min at -20°C. Samples were then centrifuged at 16,100 x g for 15 min at RT and the supernatant was removed. The pellet containing DNA and glycogen were then washed twice with 150 µL of 70% ethanol with 5 min spins at 16,100 x g. The supernatant was then removed, and the pellet was allowed to air dry in a laminar flow hood for 10 min. Finally, the dried pellet was resuspended in 20 µL of 10 mM Tris pH 8.5.

To genotype the animals, extracted DNA was amplicon sequenced using Miseq as previously described (*64*). Briefly, amplicon primers were designed to span a 465 bp and 459 bp region around guide pairs 1 and 2, respectively (table S15). One µL of genomic DNA was used in a 25 µL KOD polymerase reaction (Sigma-Aldrich, cat. no. 71086) with amplicon specific primers containing Illumina overhang adapters using the following PCR cycling parameters: one cycle of

95°C for 2 min; 35 cycles of 95°C for 20 s, 56°C for 30 s, and 70°C for 10 s; one cycle of 70°C for 10 min. The quality and size of resulting PCR products were verified with a 1% agarose gel and electrophoresis. PCR products were then purified using a Zymo Select-a-Size DNA Clean and Concentrator Magbead Kit (Zymo, cat. no. D4085) and eluted in 50 µL of 10 mM Tris pH 8.5 following manufacturer's instructions. Each PCR product was then used in a second 50 µL PCR reaction using KAPA HiFi HotStart ReadyMix (Roche Molecular Systems Inc., cat. no. 7958935001) to add dual-index barcodes for sample multiplexing and Illumina sequencing adapters (table S16), using the following PCR cycling parameters: 95°C for 3 min; 8 cycles of 95°C for 30 s, 55°C for 30 s, and 72°C for 30 s; one cycle of 72°C for 5 min. PCR product was again verified through gel electrophoresis to confirm expected quality and product size. PCR products were again purified using a Zymo Select-a-Size DNA Clean and Concentrator Magbead Kit and eluted in 30 µL of 10 mM Tris pH 8.5 following manufacturer's instructions. DNA concentration of PCR products were measured on a Nanodrop spectrophotometer, and DNA was normalized, pooled, and diluted to a final concentration of 4 nM with 25% PhiX Control v3 (Illumina no. FC-110-3001). The final library was sequenced on an Illumina MiSeq System using paired-end 250-bp reads (Illumina MiSeq Reagent Kit v3; cat. no. MS-102-3003) at Carnegie Science.

To assess mutation frequency, we used CRISPResso2 (v2.3.2). Paired-end sequences were demultiplexed and filtered for read quality using BLC Convert software (ver. 4.4.4) using default settings. The resulting sequences were aligned to reference sequences using the command *CRISPRESSOBatch* with the following parameters: quantification_window_size 2, quantification_window_center -3, min_average_read_quality 30, min_single_bp_quality 5, min_paired_end_reads_overlap 20, min_bp_quality_or_N 20, min_frequency_alleles_around_cut_to_plot 0.01, ignore_substitutions, default_min_aln_score 30. These parameters allowed *CRISPRESSOBatch* to search for mutations that begin 3 bp upstream of the start of the PAM site (the predicted cut site for S.p. Cas9) and 2 bp upstream and downstream of the cut site. These parameters also filtered reads that had a minimum average read quality of 30, a minimum single bp quality of 5, a minimum overlap of 20 bp for the paired-end reads, and a minimum alignment score to reference sequence of 30. Samples with less than 1000 reads were discarded in subsequent analyses.

*Assessing algal morphology in SLC26A11 mutants*

For Aiptasia, dark- and light-colored tentacles from SLC26A11 mutants were clipped and imaged under DIC and fluorescence (Cy5; Ex BP 620/60; Em BP 700/75) microscopy on a Leica DM6 microscope with a 40X objective. For *G. fascicularis* mutants, images described earlier for phenotyping symbiont density were used for the quantification of algal morphology. Fiji was used to quantify algal cell size and chlorophyll fluorescence intensity by randomly and manually selecting 10 algal cells from each image.

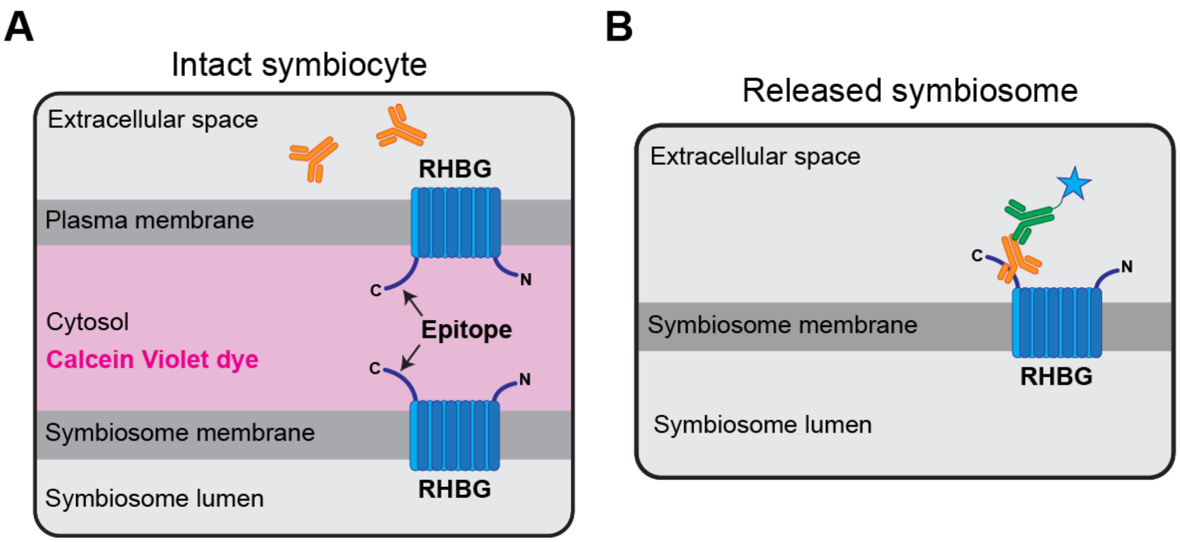


**fig. S1. Strategy to stain the symbiosome membrane of released symbiosomes with the RHBG antibody**. **(A)** The epitope for RHBG antibody is on the cytosolic side of both the symbiosome membrane and the host cell's plasma membrane. The antibody cannot bind intact symbiocytes due to the plasma membrane. **(B)** Once the symbiosome is released from a lysed symbiocyte, the antibody can bind the epitope. These properties allow for the discrimination of symbiocytes from released symbiosomes (Fig 1, B to D).

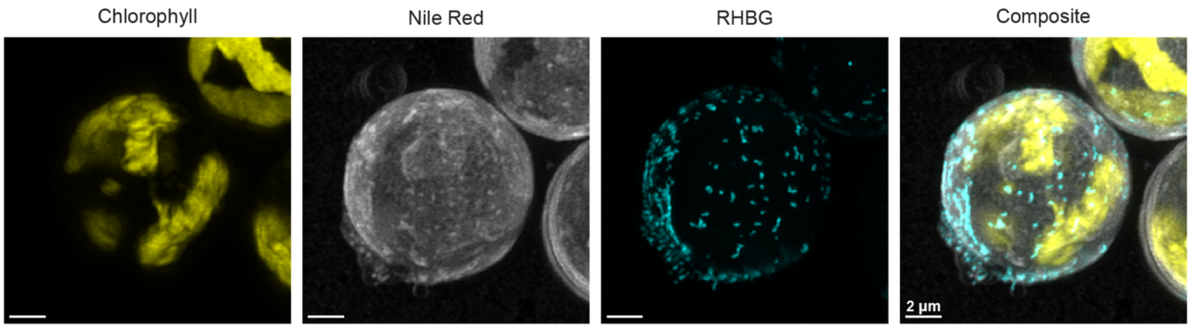


**fig. S2. RHBG localization around a released symbiosome.** A max projection of super-resolution confocal micrographs from Figure 1D and 1E shows punctate localization of RHBG protein on the symbiosome.

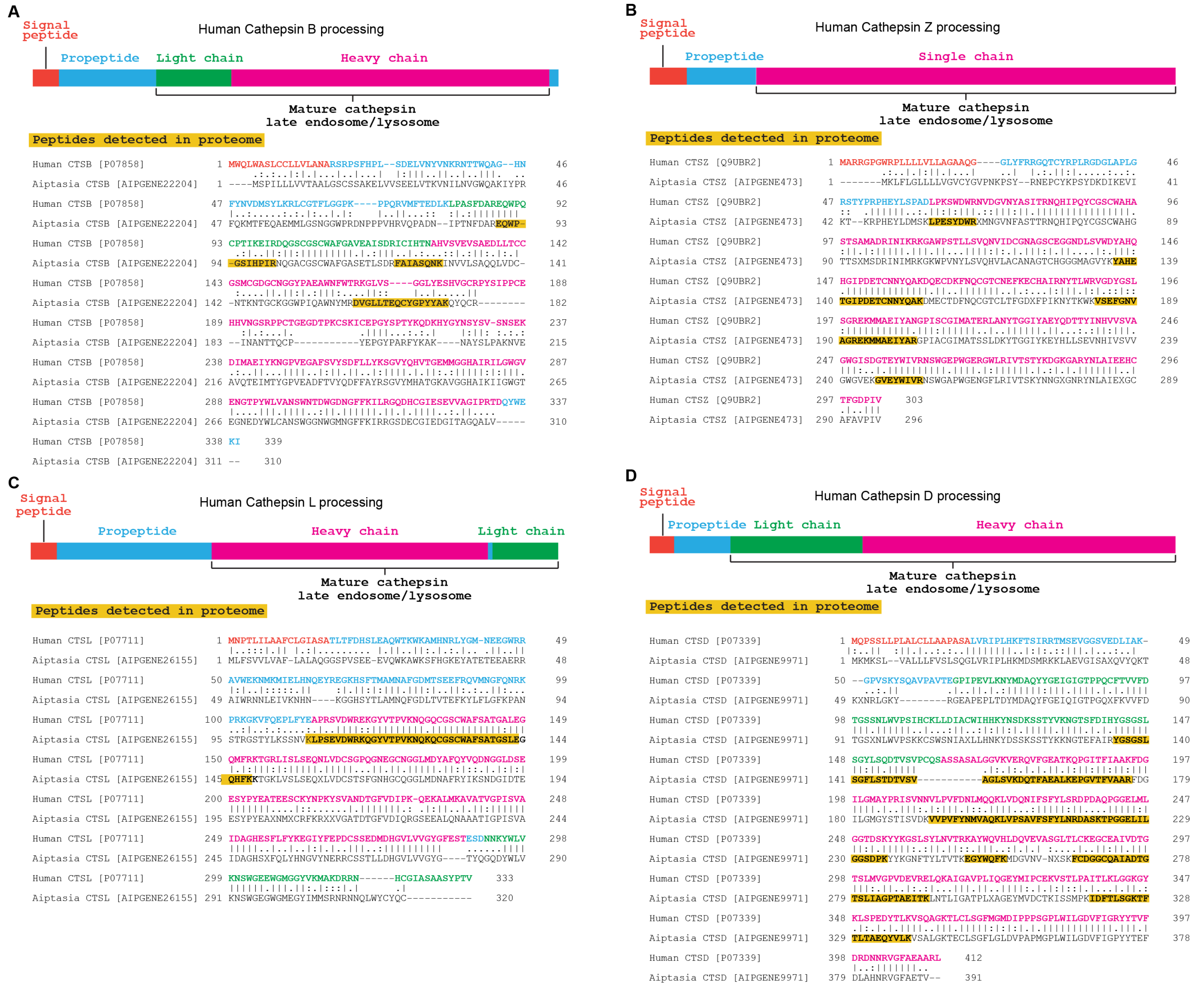


**fig. S3. Locations of peptides matching cathepsin proteins from proteome. (A-D)** Alignment of Aiptasia cathepsins proteins to their human orthologs. Colors represent different domains for each human cathepsin. Yellow highlighted regions mark positions of detected peptides for Aiptasia cathepsins from the proteome.

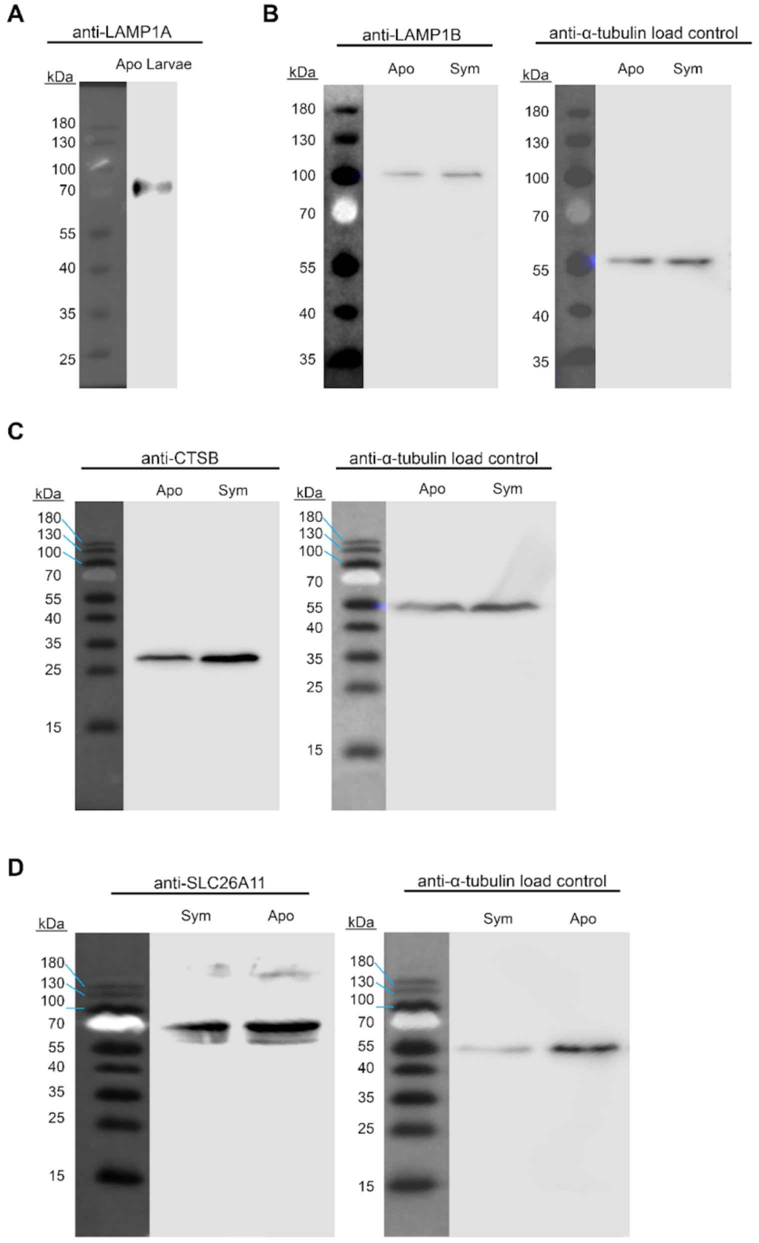


**fig. S4. Western blot confirmation of custom antibodies.** (**A**) rabbit-α-LAMP1A antibody stains a band ~70 kda in aposymbiotic larvae. (**B**) rabbit-α-LAMP1B antibody stains a band ~100 kda in size in both aposymbiotic and symbiotic polyps. (**C**) rabbit-α-CTSB antibody stains a band ~30 kda in size in both aposymbiotic and symbiotic polyps. (**D**) rabbit-α-SLC26A11 antibody strongly stains two bands that are ~55-70 kda in size in both aposymbiotic and symbiotic polyps. They are likely glycosylated and non-glycosylated forms of the protein as seen in other organisms (*65*). Mouse-α-alpha-tubulin was used as a loading control for the western blots in B-D.

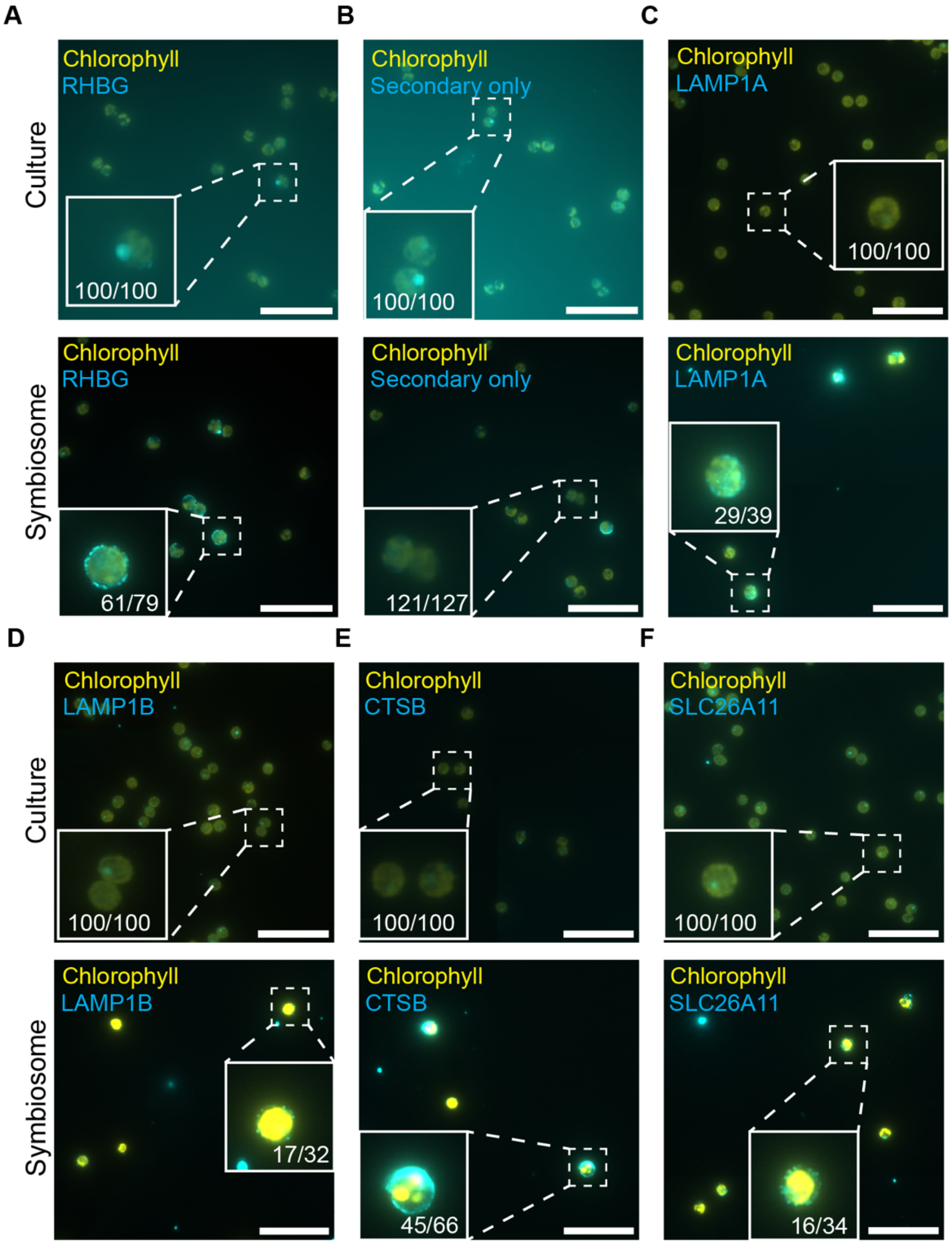


**fig. S5. Custom antibodies do not stain cultured algae. (A-F)** Staining cultured algae (top) and purified symbiosomes (bottom) with rabbit-α-RHBG, rabbit-α-LAMP1A, rabbit-α-LAMP1B, rabbit-α-CTSB, rabbit-α-SLC26A11, and Alexa Fluor™ Plus 488 goat-α-rabbit secondary antibodies show that the antibodies stained purified symbiosomes but not cultured algae. Secondary antibody only controls show lack of staining in both cultured algae and symbiosome, except for rare instances (6/127 cells) in symbiosome samples. This rare staining lacks the punctate staining that is found in all of the other antibodies. Scale bar = 5 µm.

DMSO only no lysotracker control

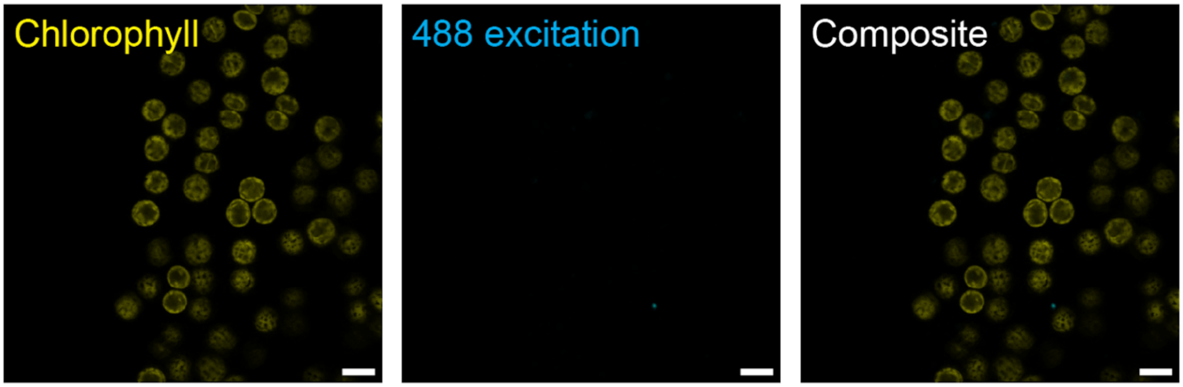


**fig. S6. DMSO carrier in lysotracker experiments does not stain symbiosomes.** Confocal image of Aiptasia tentacles treated with 0.01% DMSO does not show signal with the same settings used in Fig. 4A. Scale bar = 10 µm

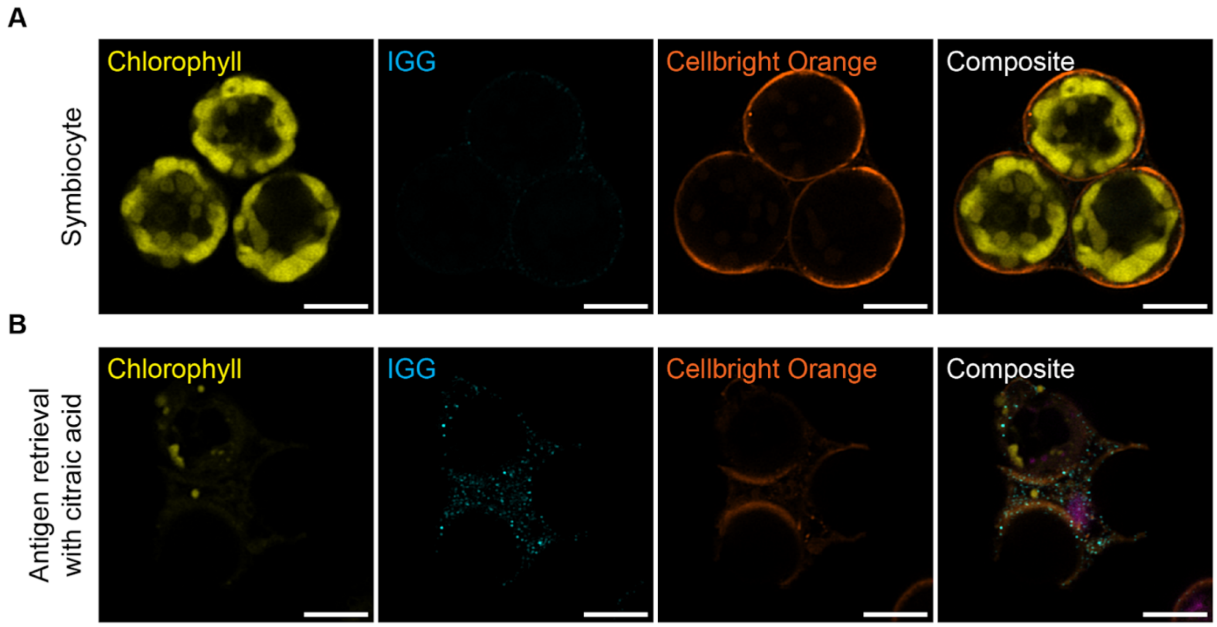


**fig. S7. Rabbit-α-IgG isotype antibody does not stain the symbiosome.** Staining of fixed symbiocytes without (**A**) and with (**B**) antigen retrieval show no or very little staining of the symbiocyte when probed with the rabbit-α-IgG antibody. This treatment serves as a control for non-specific primary antibody staining. Scale bar = 5 µm

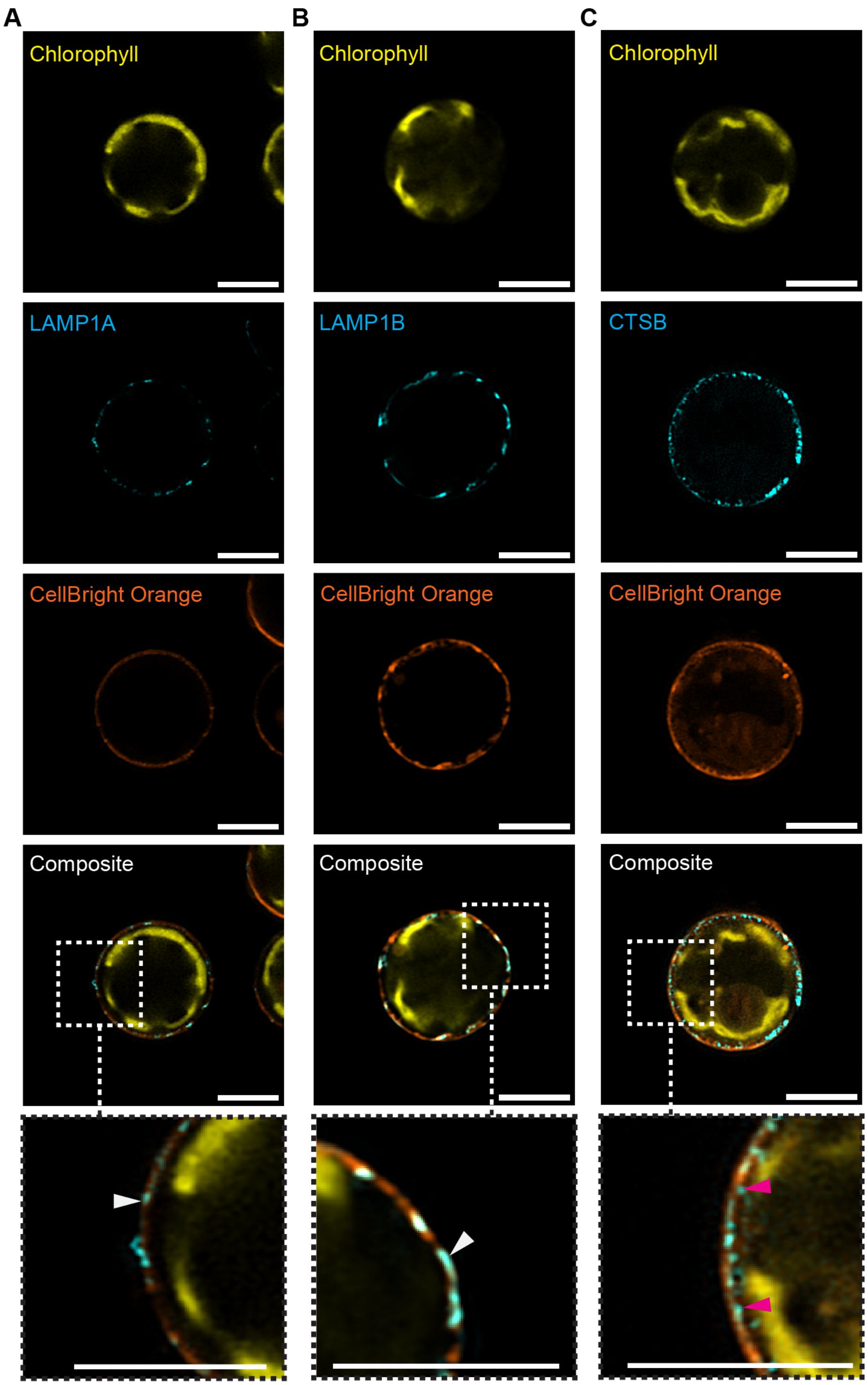


**fig. S8. Sub-symbiosome localization of LAMP1A, LAMP1B, AND CTSB proteins.**
Super-resolution confocal microscopy of immunofluorescence experiments with (**A**) LAMP1A and (**B**) LAMP1B show membrane localization on the symbiosome (white arrowheads), while (**C**) CTSB shows lumenal localization in the symbiosome (magenta arrowheads). Scale bar = 5 µm

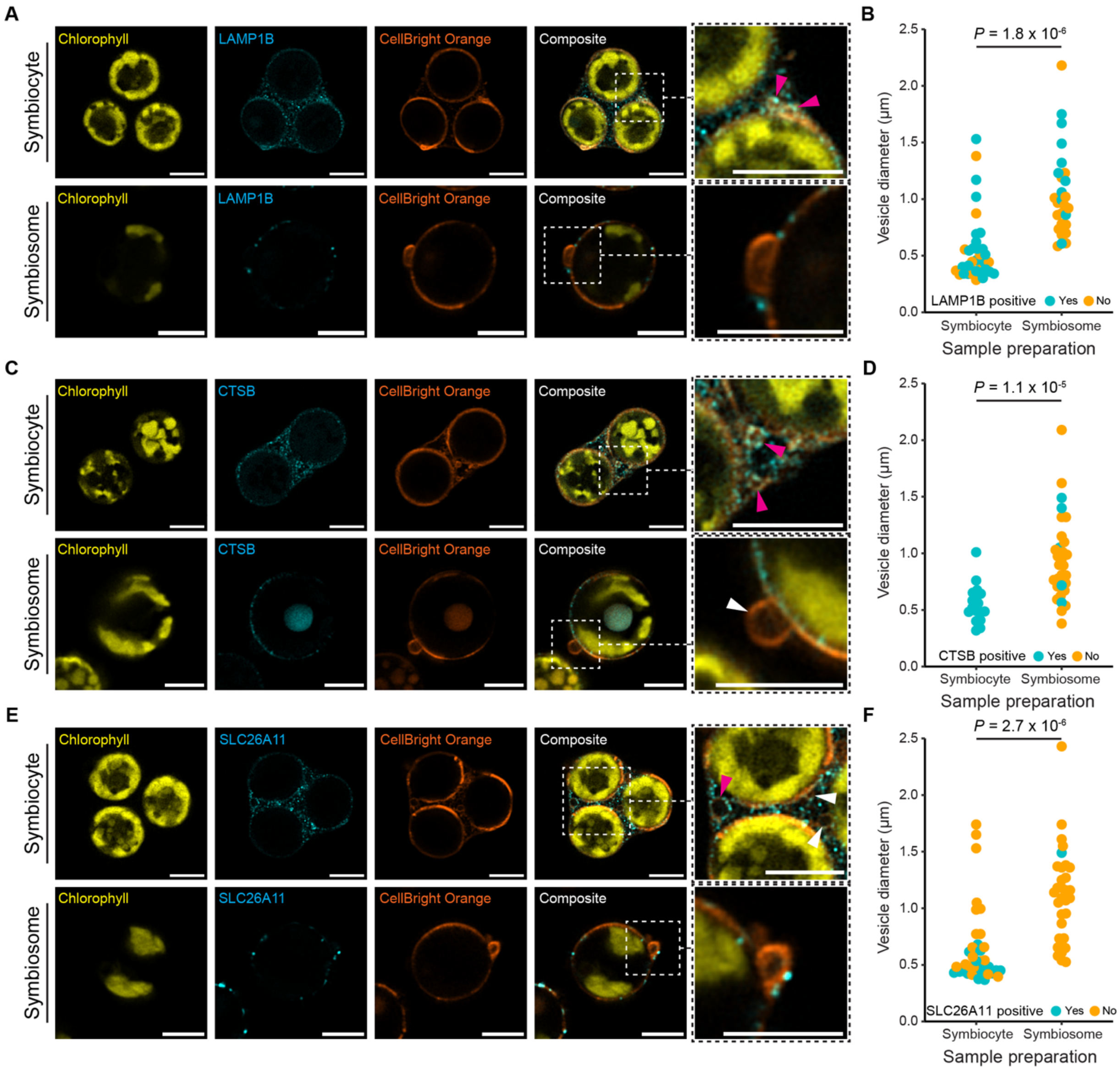


**fig. S9. LAMP1B, CTSB, and SLC26A11 proteins are localized to vesicles on symbiosomes.** (**A, C, E**) Representative super-resolution micrographs of symbiocytes and symbiosomes stained with rabbit-α-LAMP1B, rabbit-α-CTSB, or rabbit-α-SLC26A11 antibodies. Scale bar = 5 µm **(B, D, F)** Quantification of vesicle sizes that are positive or negative for the respective antibodies from symbiocytes or symbiosomes. Each dot is a vesicle and *P* values are from Welch's t-tests.

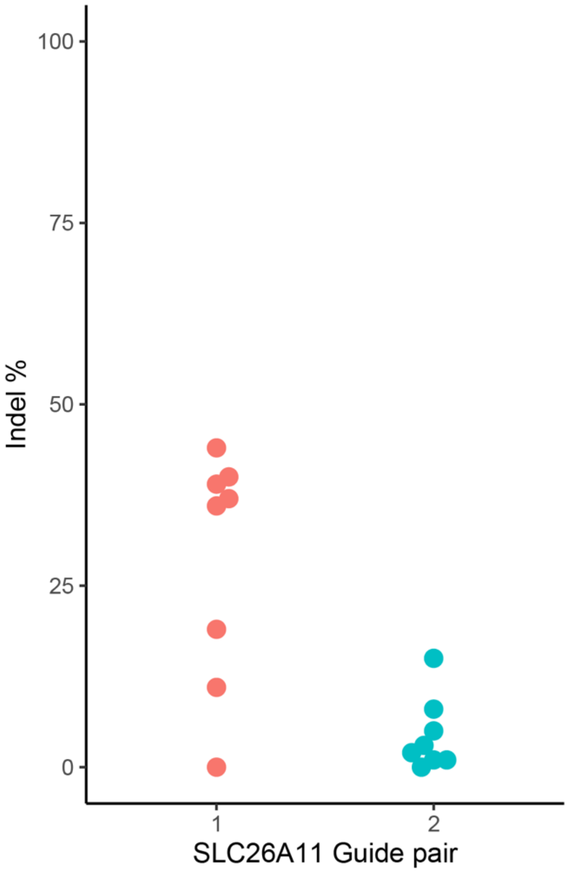


**fig. S10. CRISPR/Cas9-mediated mutations in SLC26A11 in Aiptasia larvae.** Larvae electroporated with either pair of sgRNA/Cas9 complexes were genotyped at 2 dpf. Shown are the individual larval mutation rates at each guide pair.

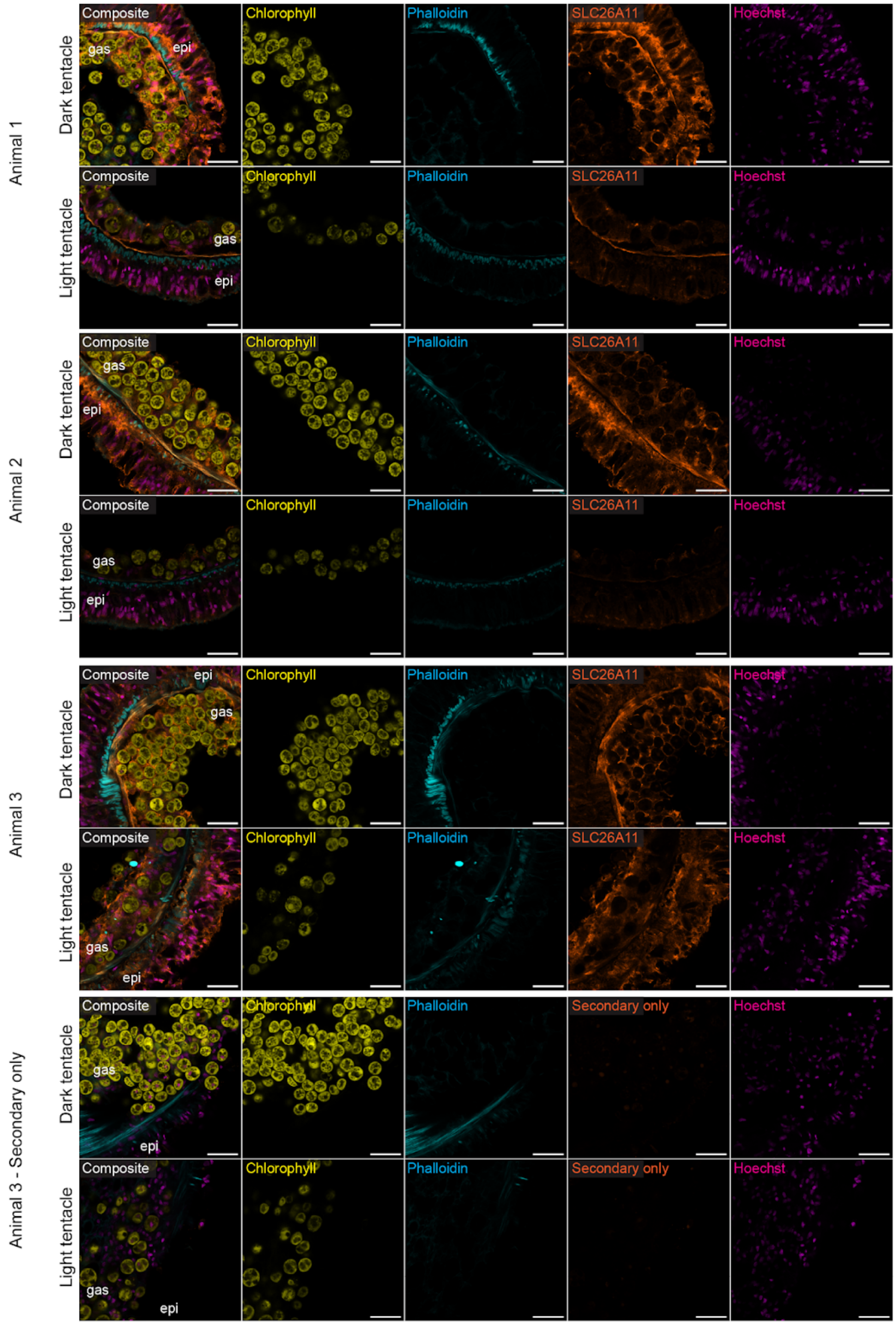


**fig. S11. Representative images of SLC26A11 localization in whole mount tentacles from three mosaic Aiptasia.** SLC26A11 was depleted from symbiosomes in light compared to dark tentacles from three mosaic animals. The secondary only controls showed no signal in either tentacle type. epi = epidermal layer, gas = gastrodermal layer. Scale bar = 20 µm

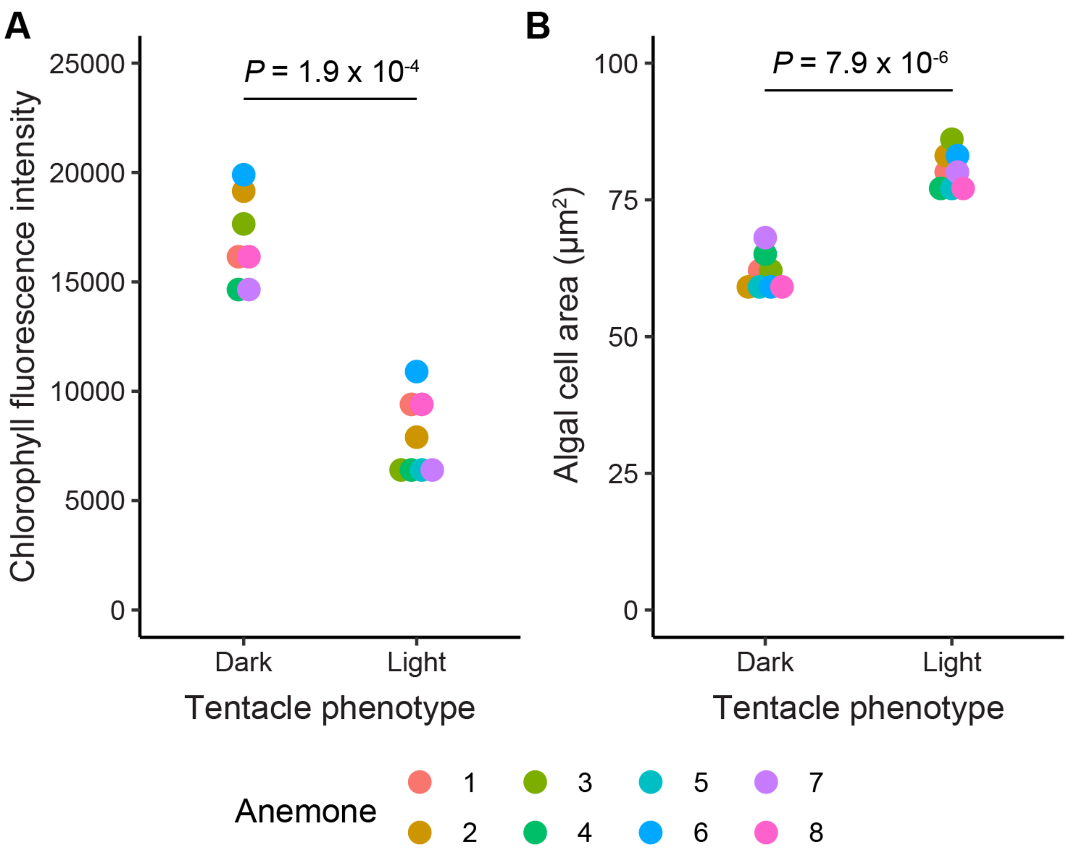


**fig. S12. Mutations in Aiptasia SLC26A11 reduces algal chlorophyll autofluorescence and increases algal size.** (**A**) Mean chlorophyll fluorescence and (**B**) mean algal cell area from paired light and dark tentacles from eight mosaic animals treated with guide pair 1. *P* values were determined by paired t-test.

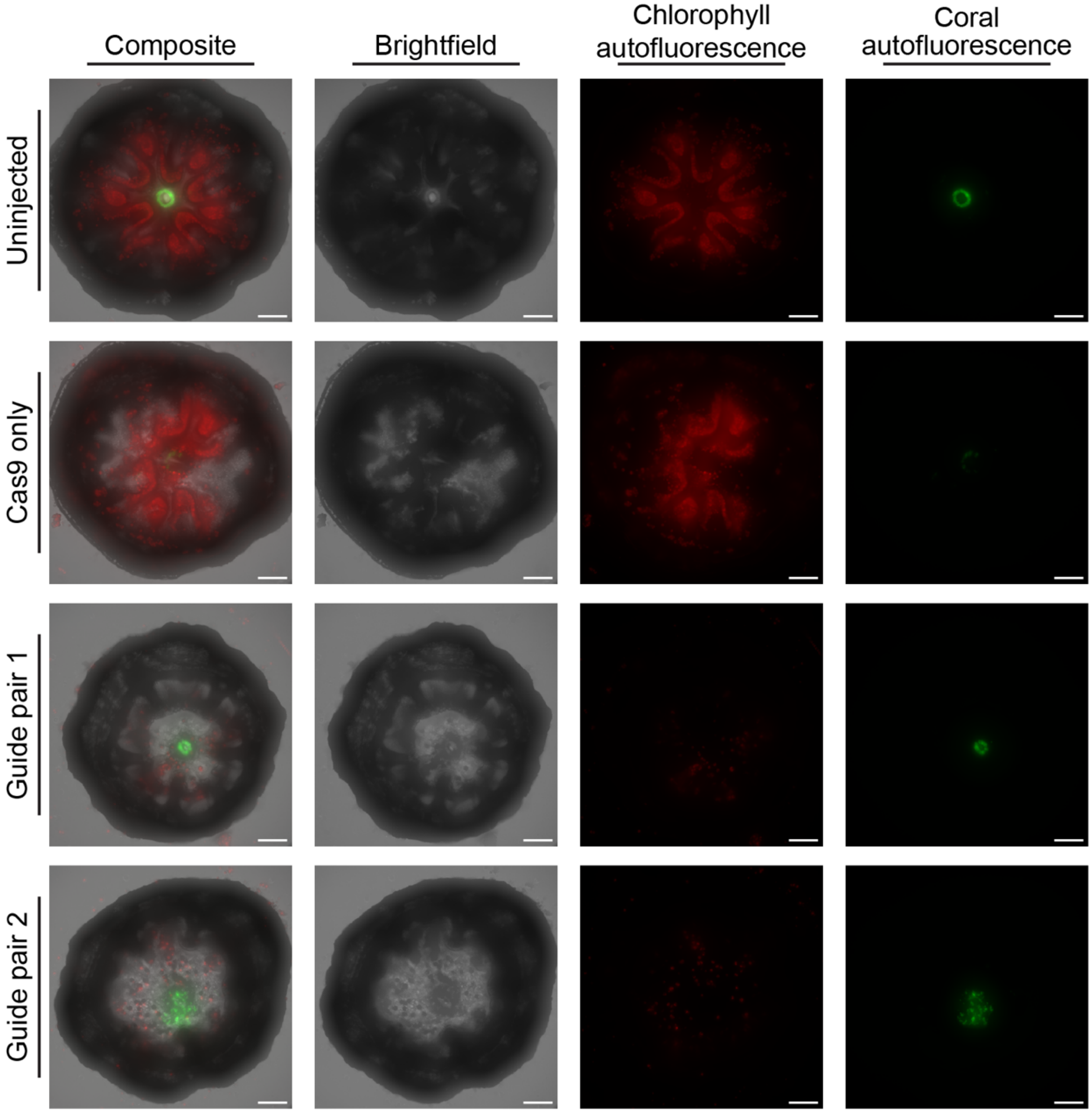


**fig. S13. *G. fascicularis* polyps from SLC26A11 CRISPR/Cas9 experiments.** Composite, brightfield, and fluorescence [chlorophyll autofluorescence (red) and animal autofluorescence (green)] micrographs of a representative uninjected animal, a Cas9-only injected animal, and animals injected with sgRNA/Cas9 complexes targeting SLC26All. These types of images were used for scoring in Fig. 5I and fig. S13. Scale bar = 100 microns.

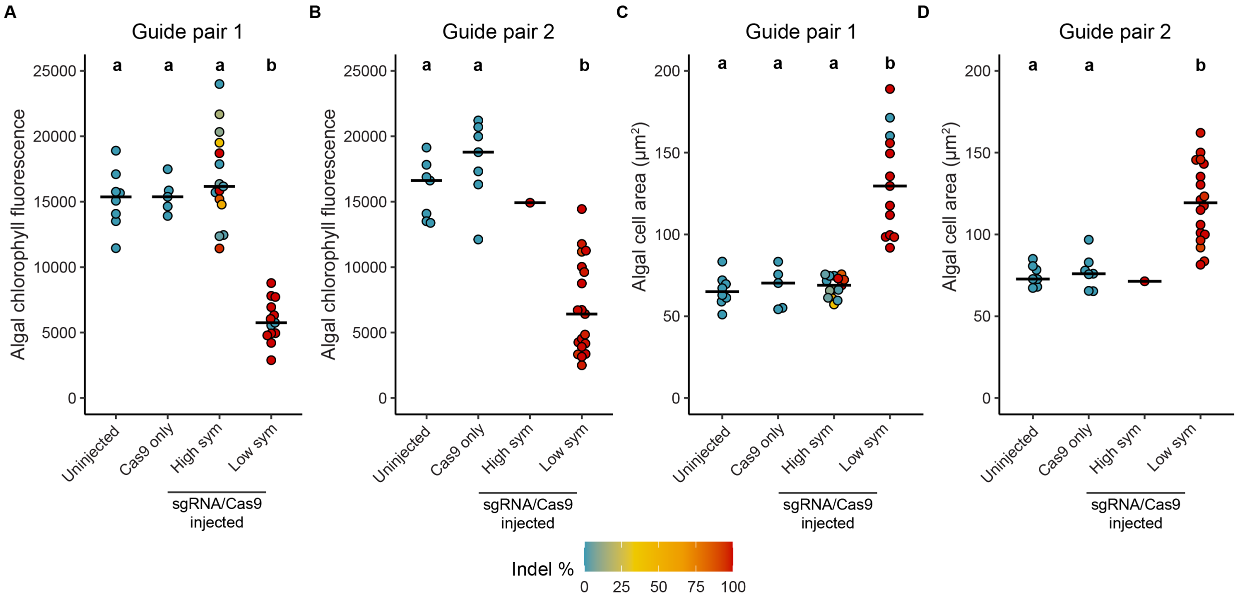


**fig. S14. Algal chlorophyll autofluorescence and cell size correlate with SLC26A11 mutation frequencies in *G. fascicularis*.** (**A,B**) Mean chlorophyll autofluorescence per alga and (**C , D**) algal cell area from *G. fascicularis* from injection experiments. The animals were separated into injection and phenotypic categories for statistical analyses. Indel percentages for each animal are shown. Letters denote significantly different groups ($P < 0.05$) determined by Kruskal-Wallis and post hoc Conover-Iman test. Significance was not determined for groups with a single animal.

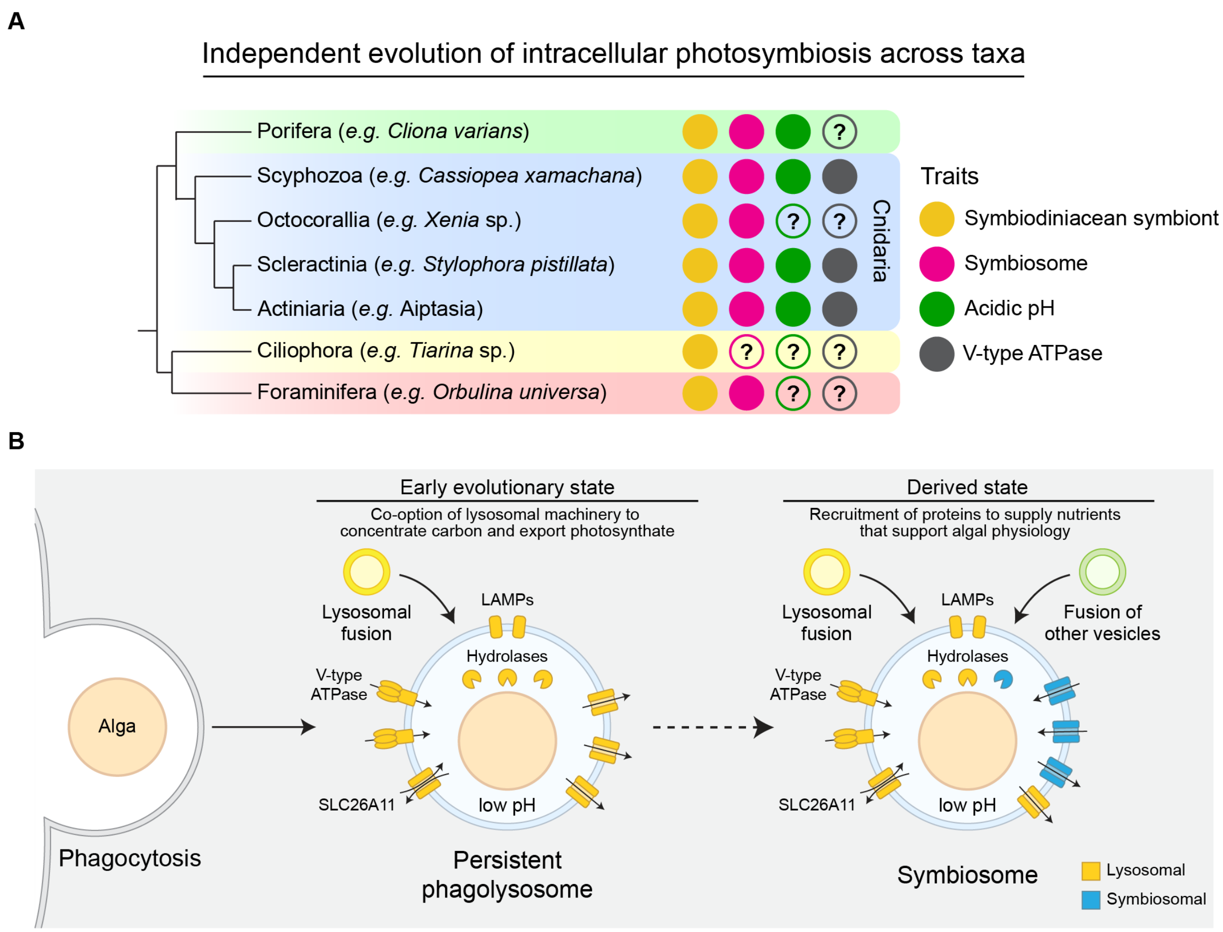


**fig. S15. Model for the repeated evolution of photosymbiosis with Symbiodiniaceae algae across phyla.** (**A**) Phylogenetic tree of organisms that have intracellular photosymbiosis with Symbiodiniaceae algae. Reported traits of symbiotic relationships are shown (*7*, *9*, *44–48*). (**B**) Proposed steps for the evolution of photosymbiosis.

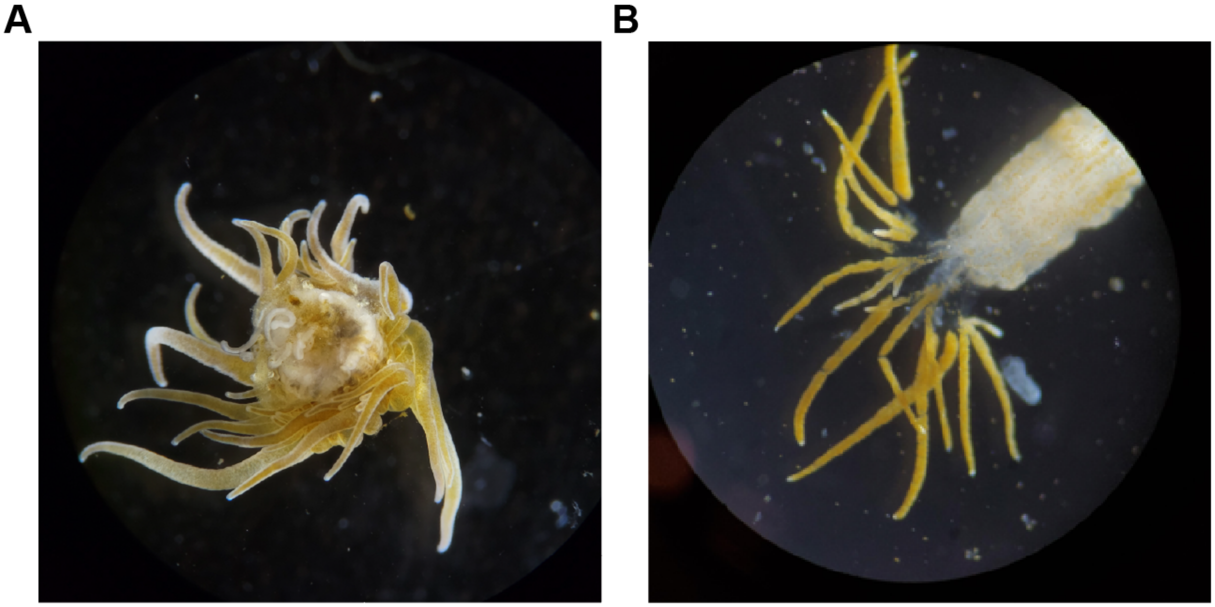


**fig. S16. Removal of epidermal tissue from anemones by incubating animals in 4% L-cysteine dissociation buffer for 1.5 h** (**A**) A symbiotic anemone before removal of epidermal tissue. (**B**) A symbiotic anemone after removal of epidermal tissue.
